# Genetic regulatory effects in response to a high cholesterol, high fat diet in baboons

**DOI:** 10.1101/2023.08.01.551489

**Authors:** Wenhe Lin, Jeffrey D. Wall, Ge Li, Deborah Newman, Yunqi Yang, Mark Abney, John L. VandeBerg, Michael Olivier, Yoav Gilad, Laura A. Cox

## Abstract

Steady-state expression quantitative trait loci (eQTLs) explain only a fraction of disease-associated loci identified through genome-wide association studies (GWAS), while eQTLs involved in gene-by-environment (GxE) interactions have rarely been characterized in humans due to experimental challenges. Using a baboon model, we found hundreds of eQTLs that emerge in adipose, liver, and muscle after prolonged exposure to high dietary fat and cholesterol. Diet-responsive eQTLs exhibit genomic localization and genic features that are distinct from steady-state eQTLs. Furthermore, the human orthologs associated with diet-responsive eQTLs are enriched for GWAS genes associated with human metabolic traits, suggesting that context-responsive eQTLs with more complex regulatory effects are likely to explain GWAS hits that do not seem to overlap with standard eQTLs. Our results highlight the complexity of genetic regulatory effects and the potential of eQTLs with disease-relevant GxE interactions in enhancing the understanding of GWAS signals for human complex disease using nonhuman primate models.

## Introduction

Genome-wide association studies (GWAS) have identified numerous non-coding loci that are associated with complex traits and diseases^1^. These loci are thought to modulate disease risk by regulating gene expression, yet only about 40-50% of non-coding GWAS hits are explained by expression quantitative trait loci (eQTLs)^2–6^. The majority of eQTLs have been discovered from healthy adult tissues in an unperturbed state, known as standard eQTLs^7^. However, eQTL effects can vary across different environmental contexts, such as tissue types^7^, cell states^8^, developmental stages^9^, and external perturbations^10^. Our ability to explain GWAS signals could be improved by expanding the scope of eQTL studies to account for gene-by-environment interactions (GxE).

Metabolic traits are highly influenced by environmental factors, particularly diet. For example, high-fat diets have been associated with dysregulation of lipid metabolism, leading to increased adiposity and the development of metabolic disorders such as obesity and type 2 diabetes^11,12^. Investigating the intricate interplay between genetics and diet is essential to advance our understanding of metabolic health and develop effective strategies for disease prevention and management.

However, studying gene-by-diet interactions in humans is challenging due to the practical limitations of performing long-term diet interventions and the restricted availability of living tissue samples. Rodent models can be used to circumvent these challenges, but due to genetic and biological differences, it is often difficult to translate results from rodents to humans, especially when GxE interactions are involved. In contrast, non-human primates share a striking genetic and physiological resemblance to humans, and their larger size permits collection of multiple tissue samples throughout the lifespan, facilitating longitudinal studies. The complex social interactions and structure of baboon colonies and a growing suite of baboon genetic resources, have made baboons one of the preferred primate models for genetic studies of complex traits^13–20^. Baboons are particularly well-suited for studying metabolic disorders, as human lipid metabolism is more strongly correlated with that of baboons than with lipid metabolism in other Old World monkeys such as rhesus macaques, cynomolgus macaques, or vervet monkeys^21^. Indeed, plasma cholesterol concentrations in baboons respond to dietary cholesterol and fat intake in a similar manner as humans^22^. In this study, we explored gene-by-diet effects on gene regulation *in vivo* in a longitudinal study of baboons.

## Results

### Modelling gene-by-diet interactions in baboons

We obtained adipose, liver, and muscle tissue samples from 99 sexually mature captive baboons (57M, 42F) on a standard low cholesterol, low fat (LCLF) diet (**Figure 1A; Table S1**). After a two-year period of feeding the baboons a high cholesterol, high fat (HCHF) diet (**Table S1**), we collected a second set of samples from the same donors and tissues (**Figure 1A**). We used RNA-sequencing to obtain transcriptomic profiles for each tissue sample. We recorded extensive metadata at every stage of sample collection and processing (**Table S2**). To confirm the tissue source of each sample, we computationally inferred cell type composition in each sample based on gene expression signatures^23^ (**Figure 1B**). Following rigorous quality control and filtering, we retained data from a total of 570 samples and 20,224 genes for downstream analyses (**Figures S1 and S2**).

**Figure 1.**
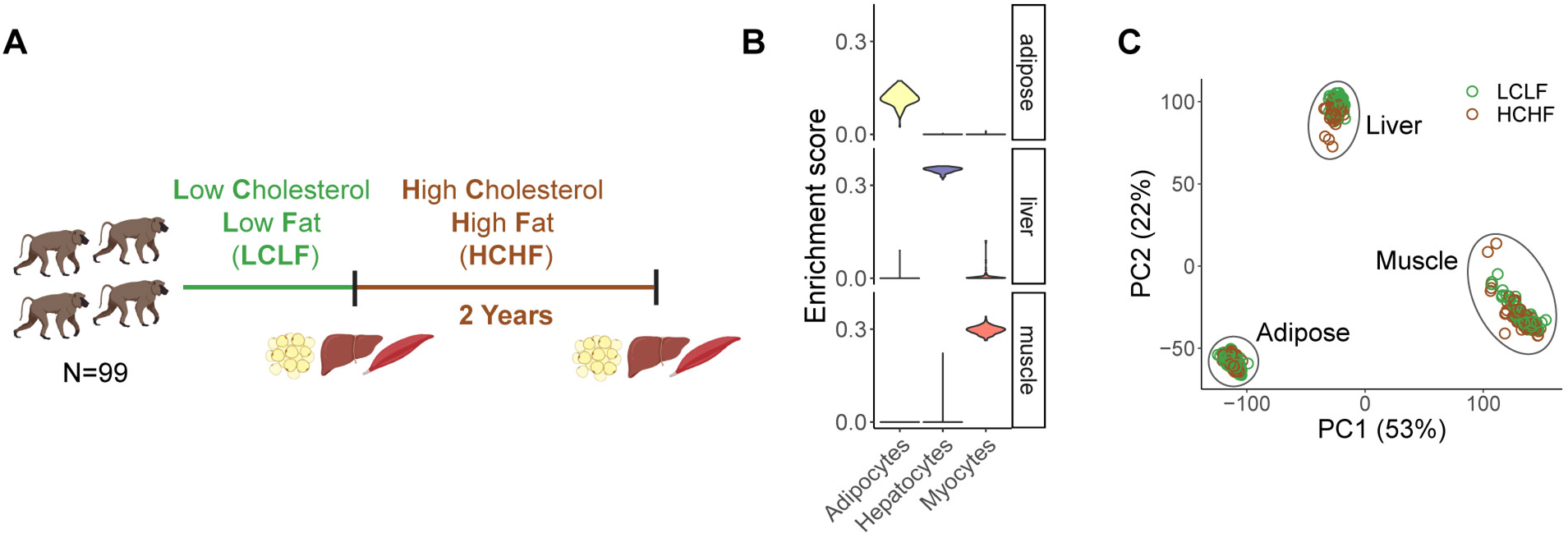
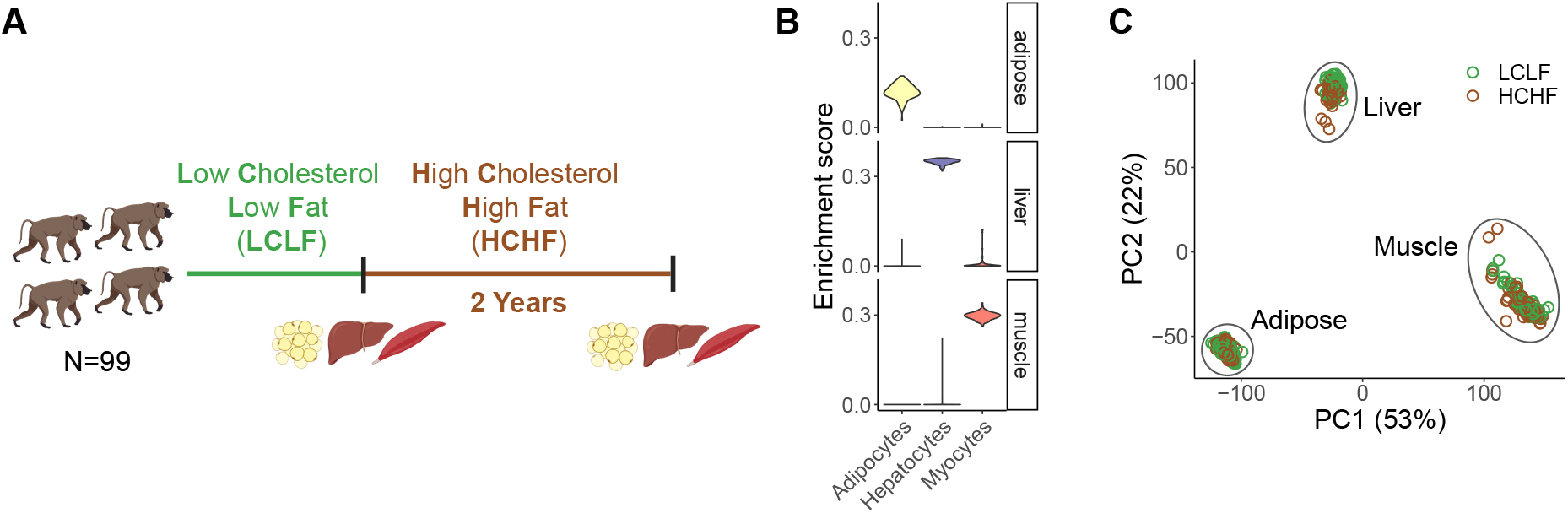
Overview of the study design and dataset. (**A**) Overview of diet intervention and sample collection. 99 baboons switched from a LCLF diet to a HCHF diet for two years. mRNA was collected from liver, muscle, and adipose tissue before and after the diet switch. Collection time points are indicated by vertical bars. (**B**) Representative cell type enrichment in each tissue. (**C**) Principal components (PC) analysis of mRNA samples after quality control (n=570), with diet condition indicated by color.

We also evaluated possible biological and technical sources of variation in the filtered data, confirming that most of the variation in the gene expression data is explained by tissue type (**Figures 1C and S1D**). Finally, we obtained genotype data for each animal from either high-coverage whole-genome sequencing (WGS) data^16^ or high-fidelity imputation^24^ from low-coverage WGS data (**Figure S3**). These data allowed us to investigate how gene expression and genetic regulation change in response to the HCHF diet in a controlled environment.

### Gene expression changes in response to HCHF diet

As a first step of our analysis, we characterized gene expression changes in response to the HCHF diet in each tissue. Across the three tissue types, we classified 6,378 genes as diet-responsive (DR); namely, genes that are differentially expressed between LCLF and HCHF conditions in at least one tissue type (false discovery rate (FDR) < 0.01, **Figure S4A; Table S3**). By jointly analyzing the DR effects, we found that more than 60% of DR genes are specific to a single tissue (**Figure S4B-C**). To understand how DR genes function at a broad level, we performed gene set enrichment analysis (GSEA) in each tissue using 50 hallmark gene sets that represent well-defined biological states or processes^25^. Among the top-scoring results, we identified positive enrichment of DR genes involved in epithelial-mesenchymal transition (EMT), inflammatory responses, interferon responses, KRAS signaling, TNF-α signaling, and other immune-related processes in adipose and liver (**Figures 2A, S5, and 6**). Both chronic inflammatory microenvironment and EMT have been shown to promote pathological fibrosis and metabolic dysfunction in tissues such as adipose and liver^26,27^.

**Figure 2.**
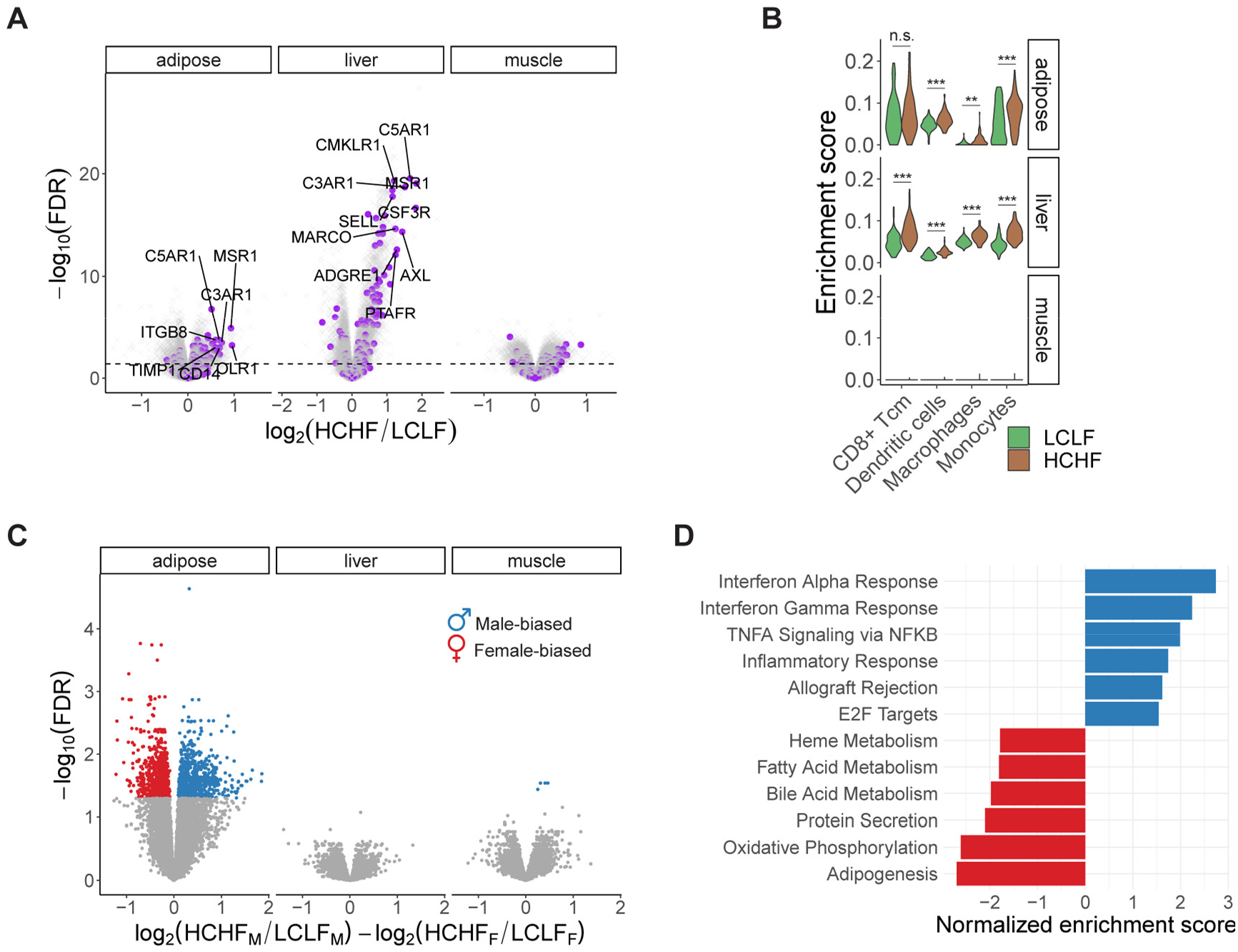
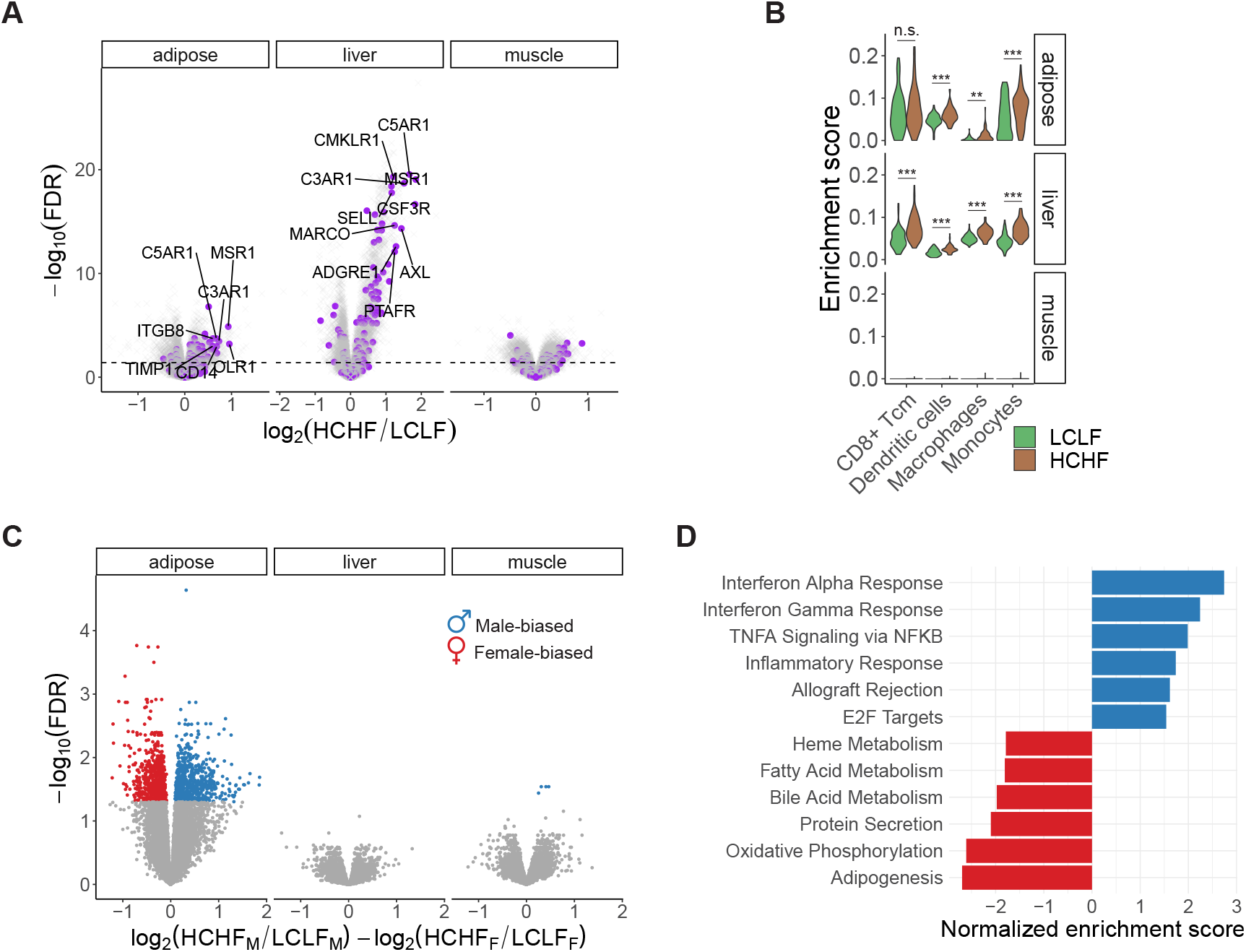
Transcriptional responses to the HCHF diet. (**A**) Diet-responsive differential expression analysis. DR genes are significantly enriched in inflammatory responses in adipose (P<2×10-10) and liver (P<6×10-16), but not in muscle (P=0.1). Inflammatory response genes are highlighted in purple. The dashed line indicates an FDR threshold of 0.05. (**B**) Cell type enrichment changes in response to the HCHF diet. Adipose and liver are significantly enriched for a variety of immune cells following the HCHF diet. Asterisks indicate statistical significance (ns: non-significance, *P<0.05, **P<0.01, ***P<0.001). (**C**) Differential expression analysis of sex-biased diet responses. Colored dots are significant sex-biased DR genes with an FDR threshold of 0.05. (**D**) Pathway enrichment of sex-biased DR genes. Top positive enrichments (FDR<0.05) are male-biased (blue) and associated with immune-related activities. Top negative enrichments (FDR<0.05) are female-biased (red), involved in metabolic processes. The enrichment items are ranked by the normalized enrichment score from top to bottom.

In rodents, excess dietary fat and cholesterol have been shown to promote an increase in tissue-resident inflammatory immune cells^28,29^. To determine whether the HCHF diet induced a similar shift in the cell composition of baboon tissues, we compared the inferred cell type composition in each tissue before and after the change in diet (**Figure S7**). We found a variety of immune cell types, including dendritic cells, monocytes, and macrophages, to be significantly more enriched following the HCHF diet, specifically in adipose and liver (**Figure 2B**). Expansion of monocyte-differentiated macrophages has been shown to promote fibrosis in human liver^27,30^. Indeed, we observed an enrichment of fibroblasts in liver after the switch to the HCHF diet (**Figure S7**).

### Sex differences in transcriptional responses to HCHF diet

Many metabolic traits show sex-differential characteristics, which could be explained by sex biases in biological processes and genetic regulation. We thus explored whether male and female baboons differed in their transcriptional responses to the HCHF diet. While we found no evidence of sex bias among DR genes in liver or muscle, we classified 1,434 sex-biased DR genes (FDR < 0.05) in adipose tissue (**Figures 2C, S8A and S8B; Table S4**). GSEA revealed that male-biased DR genes are more enriched in immune-related functions, whereas female-biased DR genes are more enriched in metabolic processes (**Figure 2D**). To explore these patterns further, we performed an independent analysis of expression data from males and females separately. Our sex-stratified analysis indicates that immune responses are enhanced in both sexes, with a stronger enrichment in males after the HCHF diet (**Figure S8C**). This may be in part due to the anti-inflammatory effects of estrogens, the primary female sex hormones^31,32^.

### Diet-responsive genetic effects on gene expression

Although we observed global transcriptional responses to HCHF diet in baboons, the dietary effects on gene expression can vary across genetic backgrounds. Captive non-human primates, even with a relatively small sample size, have exhibited their power in eQTL studies due to the homogenous environment and uniform tissue collection process^33^. Leveraging the unique advantages of the baboon model, we investigated gene-by-diet interactions and characterized diet-responsive genetic effects on gene expression. First, we mapped *cis* genetic variants (MAF > 5%) that are associated with gene expression levels (eQTLs) of their cognate genes in each tissue independently, separated by diet condition. To do this, we fit a linear model accounting for relatedness, known sample covariates, and cryptic sources of gene expression variability. Across all six combinations of tissue and diet, we discovered 7,471 genes with at least one significant *cis*-eQTL (FDR < 0.05; **Table S5**), ranging from 2,112 to 3,509 per tissue and diet combination (Figure S9A). To increase power, we also performed a joint analysis of the effects of all top variant–gene pairs (n=14,814) across all tissue and diet combinations using mashr^34^. With this more powerful approach, which leverages the information across all samples in our study, we found a total of 12,402 eQTLs (local false sign rate (LFSR) < 0.05; **Figure S9B**).

We further explored this set of eQTL-gene pairs to identify diet-responsive eQTLs (DR eQTLs) in each tissue. DR eQTLs may differ in the presence, magnitude, or direction of their effects between the different diet conditions. We identified 2,714 DR eQTL–gene pairs with diet-specific effect in at least one tissue (LFSR < 0.05, magnitude > 1.5; **Figure 3A; Table S6**). Depending on the tissue, between 49% to 58% DR eQTLs have stronger effects in the HCHF diet. We observed that the DR eQTLs are highly tissue specific. Only 6% of DR eQTLs are shared in at least two tissues, while pairwise sharing of steady-state eQTLs across tissues is 29-38% (**Figure S10**). For example, a DR eQTL (chr1:131619858:G_T; P=3.5×10^-5^) for *APOA2* emerged in response to the HCHF diet and is present only in liver (**Figure 3B**). APOA2 is involved in lipid metabolism and transport^35^, and multiple variants that are linked to *APOA2* have been found to be associated with cholesterol levels in humans^36^. An interaction between an *APOA2* single nucleotide polymorphism (SNP) and saturated fat intake has also been reported to influence body mass index (BMI) and obesity^37–39^. This non-coding SNP of *APOA2* (rs5082) shows a significant positive association with BMI in three independent cohorts of European and Hispanic individuals with high saturated fat intake (>22g/d) (**Figure 3C)**. Non-coding trait-associated loci in humans are predominantly located in open chromatin regions in relevant cell types and are enriched in gene regulatory elements, suggesting that they are mediated by altering gene regulation of nearby genes. However, no eQTLs have been found to be linked to *APOA2* in adipose, liver, or muscle from standard eQTL mapping studies such as GTEx, indicating that some eQTLs may be revealed only in specific contexts as suggested by other studies.

**Figure 3.**
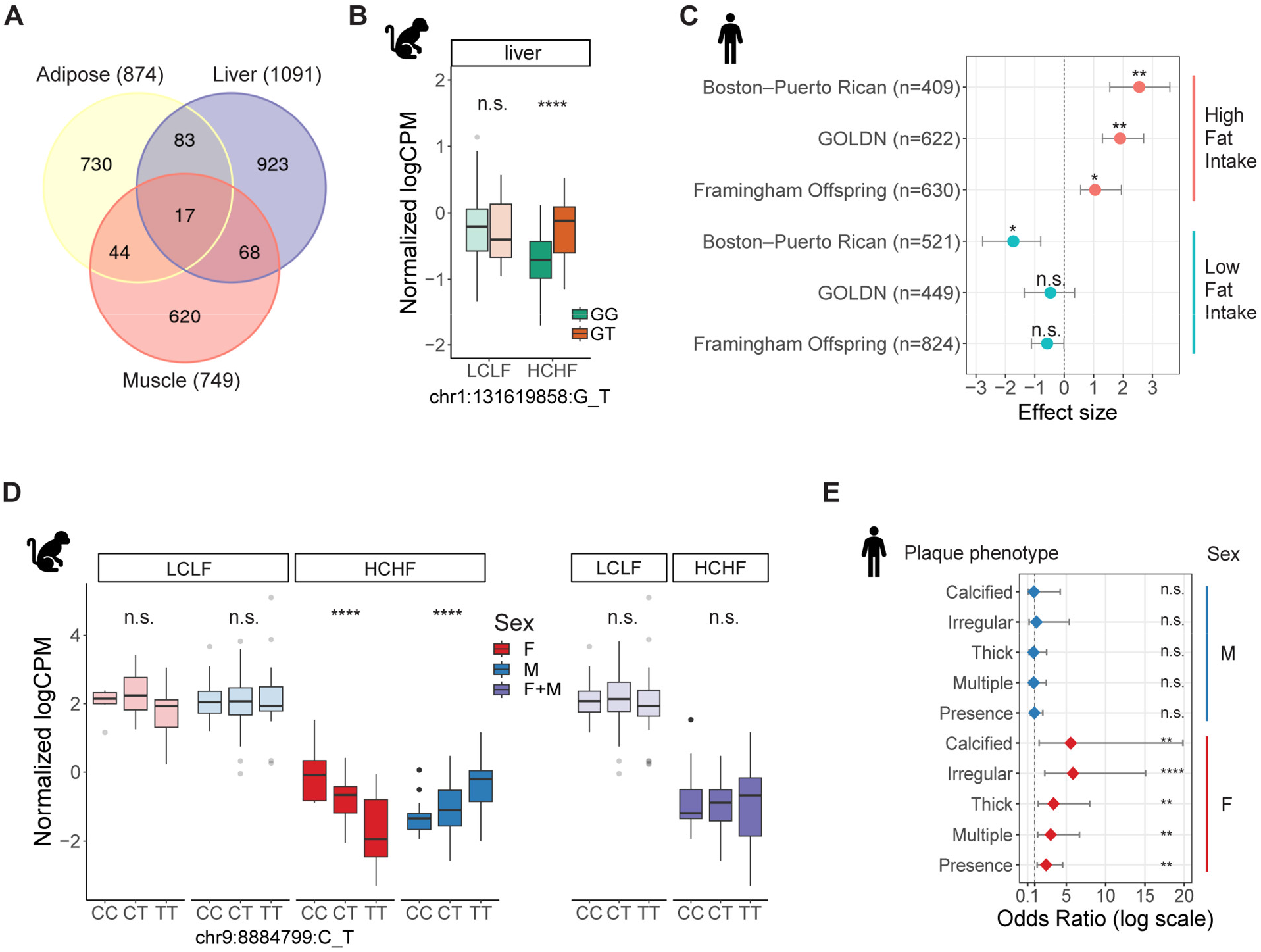
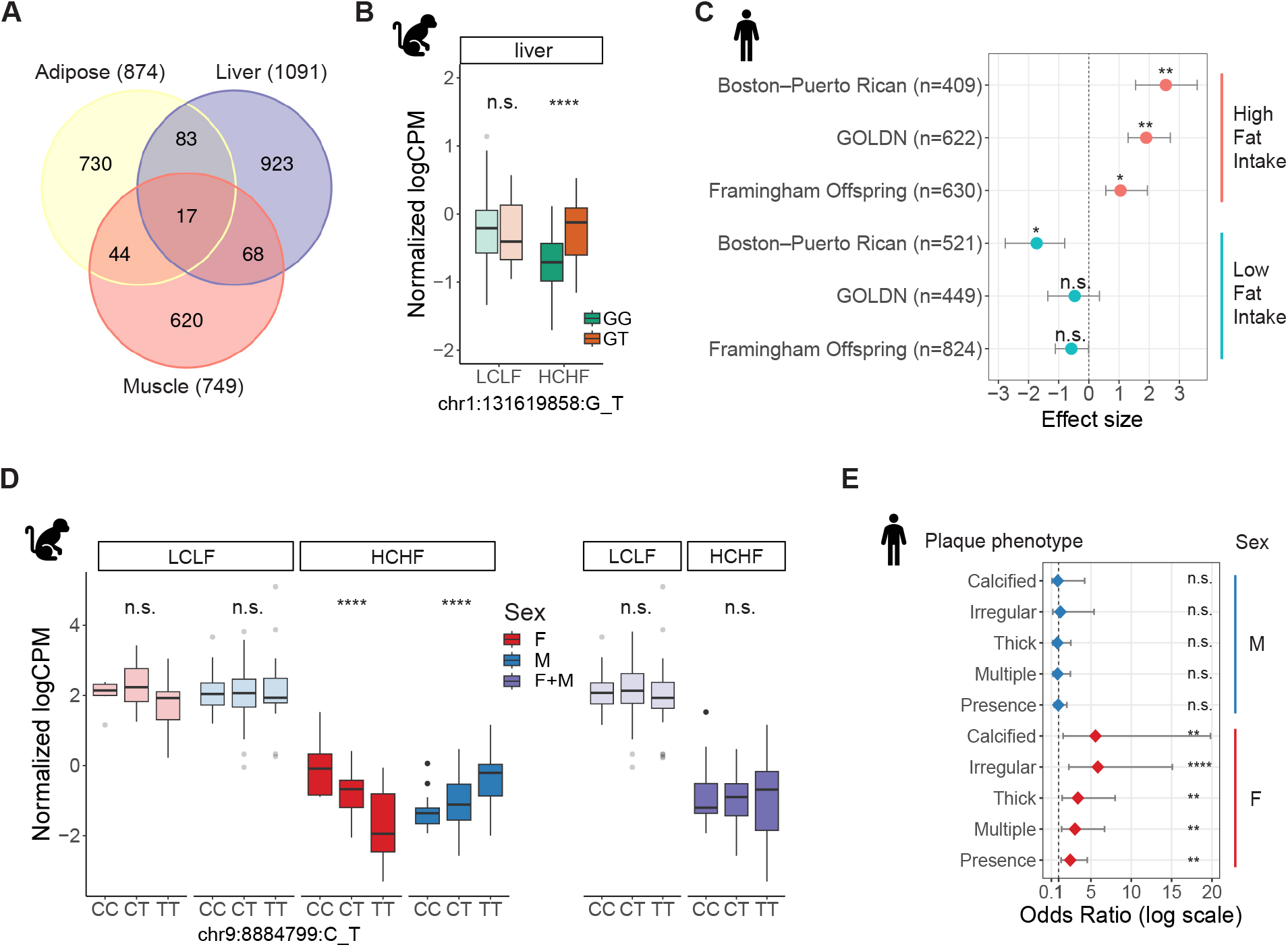
Diet-responsive genetic regulatory effects. (**A**) Discovery and overlap of DR eQTL-gene pairs across tissues. (**B**) A liver-specific DR eQTL (chr1:131619858:G_T) for APOA2. The eQTL effect emerged in response to the HCHF diet and was present only in liver (LFSRLCLF=0.49, βLCLF=0.11, LFSRHCHF=0.04, β HCHF=0.31). Opaque colors are HCHF; light colors are LCLF. Opaque colors are HCHF; light colors are LCLF. (**C**) Genetic association between a SNP of APOA2 (rs5082) and BMI from three independent human populations stratified by saturated fat intake (<22 g/d [low] and >22 g/d [high]). The populations are from the Boston–Puerto Rican Centers on Population Health and Health Disparities (Boston–Puerto Rican) Study, the Framingham Offspring Study (FOS), and the Genetics of Lipid Lowering Drugs and Diet Network (GOLDN) Study. Error bars represent standard deviation of effect size. Error bars represent standard deviations. (**D**) A sex-biased eQTL (chr9:8884799:C_T) for OLR1. The regulatory effect of this SNP was present in both males (blue) and females (red) with a discordant allelic effect, but it was not detected in sex-combined analysis (purple). Opaque colors are HCHF; light colors are LCLF. Opaque colors are HCHF; light colors are LCLF. (**E**) Sex-specific association between the variant in OLR1 (rs11053646) and carotid plaque from 287 Dominican Hispanic individuals in the Genetic Determinants of Subclinical Carotid Disease Study. The phenotypes include plaque presence and sub-phenotypes (multiple, thick, irregular, and calcified plaque), determined by high resolution B-mode carotid ultrasound. Asterisks indicate statistical significance (ns: non-significance, *P<0.05, **P<0.01, ***P<0.001, ****P<0.0001). Error bars represent 95% confidence interval of odds ratio. Error bars correspond to 95% confidence intervals.

### Colocalization of diet-responsive eQTLs with lipid biomarkers

Next, we sought to assess whether DR eQTLs can explain the molecular basis of relevant physiological traits. We performed physiological QTL mapping for 10 lipid biomarkers collected from the same cohort of baboons before and after the HCHF diet intervention^40^ (**Table S7, Figure S11**). Using a multi-trait colocalization method^41^, we identified seven regions where an eQTL colocalized with at least one lipid biomarker (posterior probability of sharing the same causal variant (PP) > 0.5; **Figure S12**). All seven colocalizations are linked to genes that contain variants associated with metabolic-relevant traits in human GWAS. Within these seven regions, we found two DR eQTL-lipid biomarker colocalizations. For example, DR eQTL (chr2:74238226:C_T) for *MAGI1* colocalized with a variant associated with the size distribution of high-density lipoprotein (dHDL) in baboons (PP=0.54, PP_candidate_SNP_=0.91, **Figure S13A**). *MAGI1* is genetically associated with a variety of relevant human traits, including cholesteryl ester (17:0) levels^42^ and coronary artery calcification^43^. The two DR eQTL–lipid biomarker colocalizations we identified have similar effects on the corresponding lipid biomarkers in both diet conditions (**Figure S13**), suggesting that the mediated effects of eQTLs on phenotypes can be context-dependent. Despite the limited sample size, our results suggest that DR eQTLs, and more generally, dynamic eQTLs identified in disease-relevant contexts, are useful for characterizing GWAS loci in humans.

### Sex biases in diet-responsive genetic effects

Given that we observed sex-biased transcriptional responses to the HCHF diet in adipose tissue, we sought to investigate sex effects on diet-responsive gene regulation. We mapped sex-biased eQTLs by incorporating a genotype-by-sex (G x Sex) interaction term into our models for each tissue and diet combination. Across all tissues, we found eight sexually dimorphic DR eQTLs (FDR < 0.25) (**Figure S14**), which are either present exclusively in one sex, present in both sexes with a concordant allelic effect but different effect sizes, or present in both sexes with a discordant allelic effect. More than half of sex-biased eQTLs are observed only in response to diet, indicating that sex effects can mask diet-responsive genetic effects (**Figures S14A-E**).

For example, we found a DR eQTL for *OLR1* (chr9:8884799:C_T) in muscle has opposite effects in males and females and was not discovered in our sex-combined analyses (P_GxSex_=1.5×10^-7^; **Figure 3D**). *OLR1* encodes lectin-like oxidized low-density lipoprotein receptor and has been shown to be associated with its protein level in GWAS^44^. It also harbors a missense variant (rs11053646; K167N) in humans that has been reported to have a sex-specific association with carotid atherosclerotic plaque in individuals of Dominican descent^45^. Sex-stratified analysis revealed that rs11053646 is significantly associated with plaque presence and all plaque sub-phenotypes in women (OR, 2.44 to 5.86; P=3×10^-4^ to 8×10^-3^) but not in men (OR, 0.85 to 1.22; P=0.77 to 0.92; **Figure 3E**).

### Genetic architecture of diet-responsive gene expression

It is recognized that eQTLs tend to exhibit a strong enrichment near transcription start sites (TSS)^46–48^. The eQTLs identified in this study follow the same enrichment pattern when considered together (**Figure 4A**). However, when we examine non-DR eQTLs (i.e. eQTLs with similar effects between the diets in a given tissue) and DR eQTLs separately, a distinct localization pattern emerges, with DR eQTLs showing only a modest enrichment near TSS (**Figure 4A**). The genomic location of an eQTL generally reflects the underlying regulatory landscape and functional activity of the target gene^49–51^. To investigate the genetic architecture of diet-responsive gene expression and determine how it differs from that of standard non-responsive gene expression, we analyzed the target genes that are associated with DR eQTLs (DR eGenes), steady-state non-DR eQTLs (non-DR eGenes), and any eQTLs (all eGenes) in each tissue. We assessed regulatory complexity and evidence of selective constraint for each group of eGenes.

**Figure 4.**
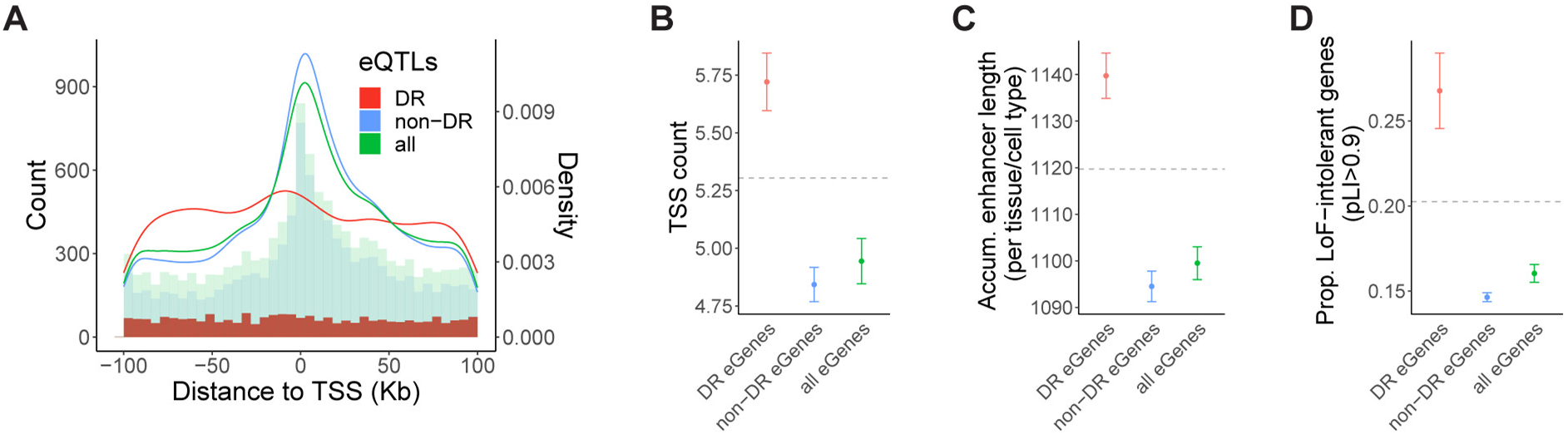
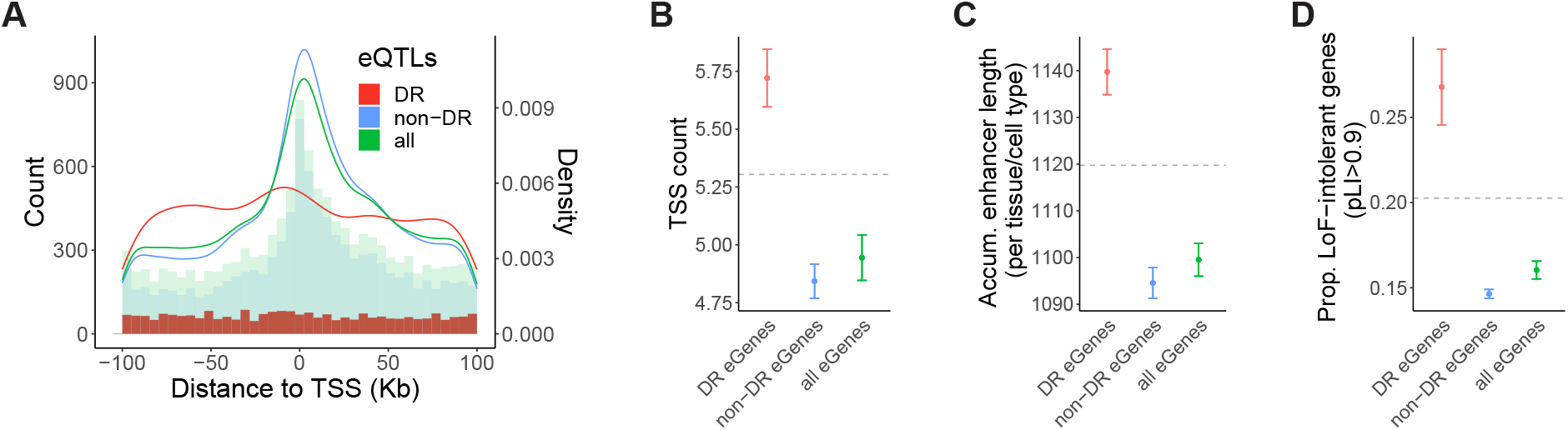
Genetic architecture of diet-responsive gene expression. (**A**) Distance of DR eQTLs (red), nonDR eQTLs (blue), and all eQTLs (green) to the TSS of the target gene. The overlaid histograms show distance-to-TSS distribution in each group of eQTLs in 5Kb bins. The curves show the kernel density estimate for each group of eQTLs. (**B-D**) Genic features of DR eGenes (red), nonDR eGenes (blue), and all eGenes (green), respectively. Horizontal dashed lines depict the average number of the corresponding feature across all genes that were tested in eQTL mapping. (**B**)Average number of TSS. (**C**) Average accumulative enhancer length per tissue/cell type. (**D**) Fraction of selectively constrained genes (pLI > 0.9). Error bars represent standard deviations.

A gene can initiate transcription from multiple TSS^48^. The regulation of TSS selection plays a crucial role in gene expression diversity and complexity, enabling context-specific transcriptional programs^52^. To explore the potential association between diet responsiveness and TSS usage, we computed the number of TSSs that are used for a given gene across a diverse set of cell types using publicly available summary statistics from the FANTOM project^50^. Across all genes, the average number of TSS is 5.3. However, DR eGenes have a higher average TSS count of 5.7, compared to an average of 4.8 TSSs in non-DR eGenes (P=6.4×10^-4^; **Figure 4B**).

Enhancers facilitate precise spatiotemporal control of gene expression by integrating signals from various cellular and environmental cues^53–55^. To assess eGene enhancer activity, we calculated the accumulative enhancer length for a given eGene across 131 tissue or cell types using enhancer-gene predictions from an activity-by-contact model^56^. Our analysis revealed that DR eGenes have a longer accumulative enhancer length per active tissue/cell type compared to the average gene (**Figure 4C**). In contrast, non-DR eGenes had a significantly shorter accumulative enhancer length than DR eGenes (P=2×10^-4^; **Figure 4C**), suggesting the diet-responsive gene regulation may be in part modulated through enhancer activity.

Previous evidence suggests that natural selection plays a significant role in shaping the genetic architecture of complex traits^57,58^. It has been observed that selectively constrained genes are depleted in eQTL genes^59^. Consistent with previous findings, we found that genes depleted of loss-of-function (LoF) variants, as measured by the pLI score, are underrepresented in all eGenes and non-DR eGenes (**Figure 4D**). However, DR eGenes display an enrichment in LoF-intolerant genes, showing a significantly higher proportion of high-pLI genes compared to non-DR eGenes (P=5×10^-3^; **Figure 4D**). We also observed that DR eGenes have overall higher minor allele frequencies (MAF) than non-DR eGenes, despite the fact that stronger selective pressure is often associated with lower MAF (**Figure S15**). This might be explained by that higher MAF gives more power to detect eQTLs with variable effects across multiple contexts as they frequently have smaller effect sizes than standard eQTLs that are shared across conditions^60^.

### Functional features of diet-responsive eGenes

The distinct regulatory architectures and natural selection signatures we observed for DR and non-DR eGenes could reflect functional differences. To investigate the functional features of DR eGenes, we performed gene enrichment for a list of broadly defined Gene Ontology (GO) biological process terms. Many functional categories show clear depletion of eQTLs, with the depletion being more pronounced in non-DR eGenes (**Figure S16A**). The depletion is more prominent for categories with a higher average pLI such as processes related to transcriptional regulation (“positive regulation of transcription by RNA polymerase II” and “negative regulation of transcription, DNA-templated”). This pattern suggests that selection purges eQTLs for genes with evolutionarily important functions. In contrast, a variety of GO categories are enriched among DR eGenes. This pervasive enrichment has also been observed for GWAS genes^6^, suggesting a functional similarity between DR eGenes and genes that are associated with complex traits.

Many biological functions are modulated by transcription factors (TFs), which play a crucial role in regulating gene expression, cell fate, development, and responses to environmental stimuli^61^. TFs have consistently been observed to be underrepresented among eGenes. To examine whether DR eGenes differ in TF representation, we calculated the proportion of TFs in each category of eGenes. We found a significant depletion of TFs in non-DR eGenes but a slight enrichment in DR eGenes (**Figure S16B**). This observation suggests that TFs may play a more prominent role in mediating diet-responsive gene expression, potentially contributing to their functional significance in the context of dietary influences.

### Enrichment of baboon diet-responsive eGenes in human GWAS

We have identified several features that distinguish DR eQTLs from standard eQTLs, including a suggestive functional resemblance between DR eGenes and GWAS genes. To explore whether diet-responsive genetic effects in baboons can provide functional insight into human non-coding GWAS variants, we integrated our results with results from human GWAS. Although SNPs are typically not conserved across species^62^, the human orthologs of eQTL genes identified in other primates tend to be associated with eQTLs in humans^33,63^. Consistently, we observed significant enrichment of baboon eGenes with a known human ortholog among human eGenes from GTEx, particularly in matched tissue types (**Figure S17**). Therefore, we focused on comparing baboon DR eGenes identified in our study and human GWAS genes from public datasets.

We tested for enrichment of baboon DR eGenes in human GWAS genes for 22 HCHF-related traits using publicly available summary statistics from UK Biobank^64^. We found that the human orthologs of DR eGenes identified in baboons are enriched among GWAS genes for most of the HCHF-related traits (**Figure 5A**). Notable enrichments include BMI, cholesterol traits, obesity, peripheral atherosclerosis, and liver cirrhosis, which are relevant to dietary fat or cholesterol intake, and are manifested in the tissue types collected in this study. However, non-DR eGenes show consistent depletion in all HCHF-related traits except for coronary atherosclerosis (**Figure 5A**).

**Figure 5.**
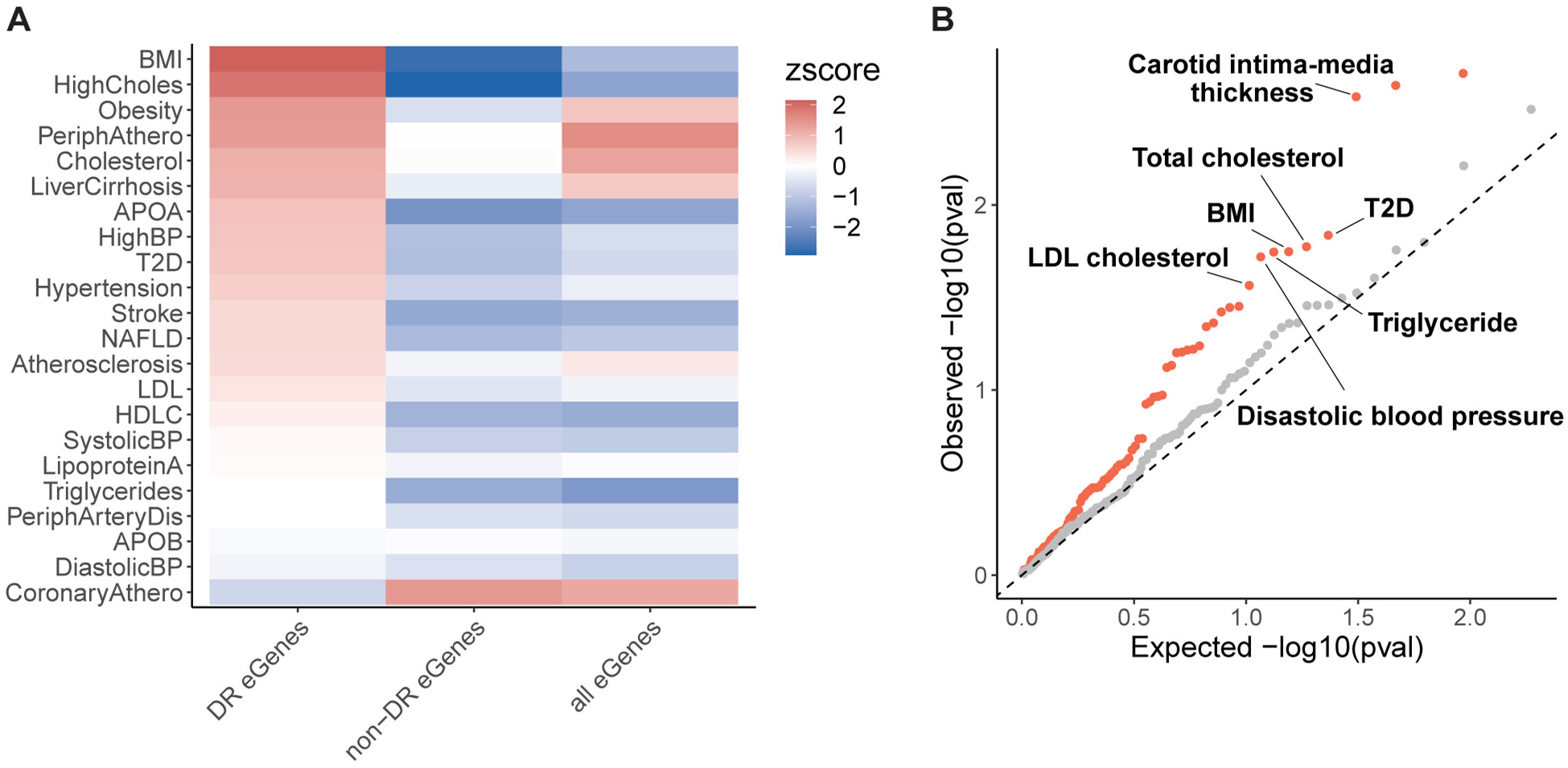
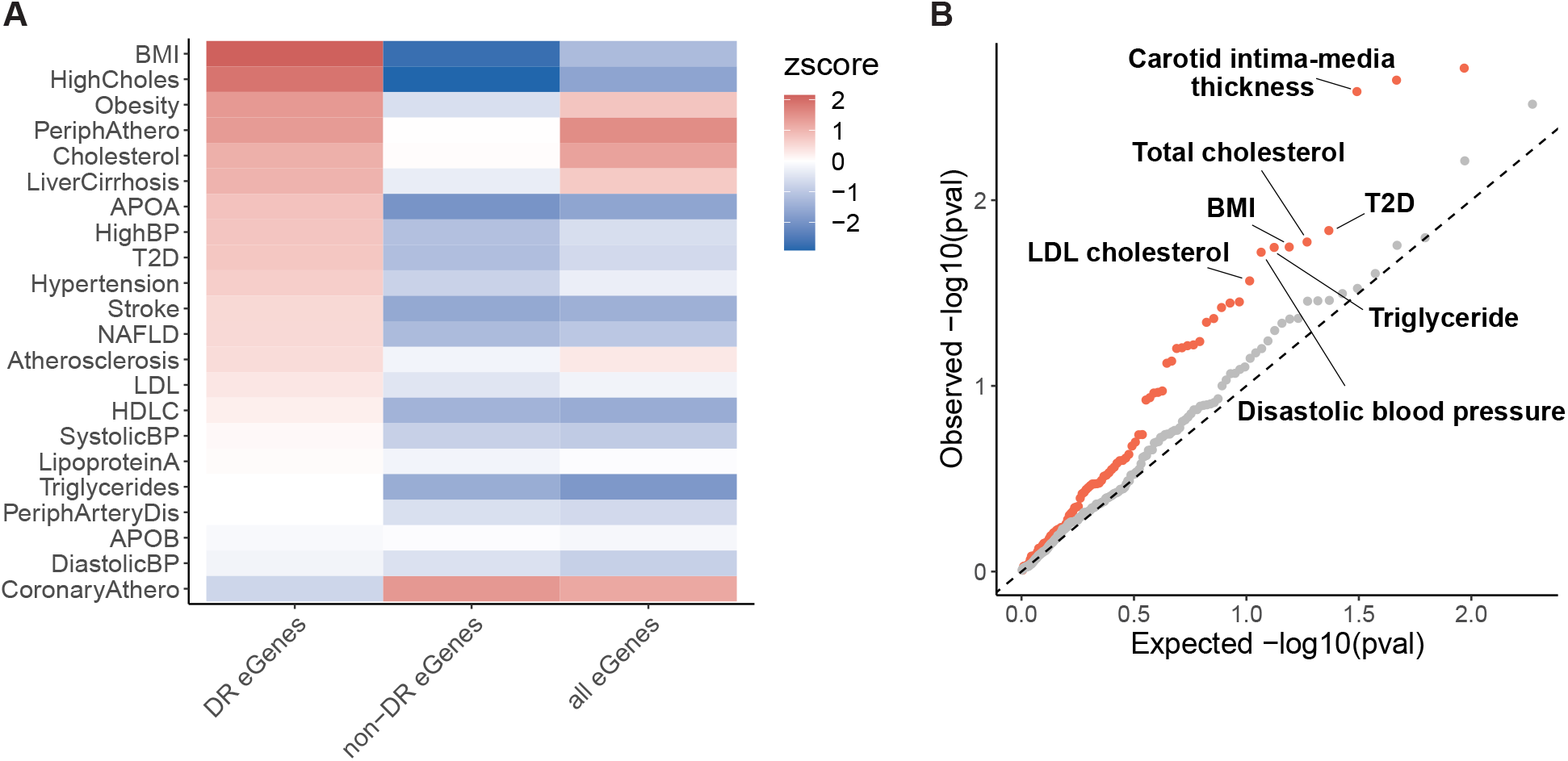
Enrichment of baboon DR eGenes in human GWAS. (**A**) Enrichment of human GWAS genes for 22 HCHF-related traits among DR eGenes, non-DR eGenes, and all eGenes. The color map represents enrichment (red) or depletion (blue) by enrichment Z scores. (**B**) Q-Q plot for enrichment of baboon DR eGenes in human trait- or disease-associated genes from GWAS. Orange dots correspond to p-values of enrichment tests for metabolic-relevant traits relative to uniformly distributed p-values (dashed line). Gray dots correspond to p-values of enrichment tests for non-metabolic traits.

To assess the specificity of the representation of DR eGenes in GWAS, we expanded our analysis to more traits and diseases using publicly available summary statistics from the NHGRI-EBI GWAS Catalog^65^, which encompasses all published GWAS. We assessed the relevance of diet to a broad range of traits, including 98 metabolic traits and 213 non-metabolic traits. We performed enrichment tests for DR eGenes in GWAS genes for each trait. Our results revealed that DR eGenes are systematically enriched in genes associated with metabolic-relevant traits (P_combined_=3.8×10^-5^; **Table S8**), with prominent signals in traits relevant to a HCHF diet, such as carotid intima-media thickness, total cholesterol, LDL cholesterol, BMI, and triglyceride levels (**Figure 5B**). When we performed the same analysis for non-DR eGenes with similar expression levels or genes that are differentially expressed in response to the diet, we observed no enrichment, suggesting that this enrichment was not confounded by differences in gene expression levels or differential transcriptional response to diet (**Figures S18A and S18B**). In contrast, we observed no enrichment of DR eGenes in non-metabolic traits (P_combined_=0.12; **Figure 5B; Table S9**). In fact, we observed a depletion of non-DR eGenes in both groups of traits (**Figure S18C**), which is consistent with our previous observation that steady-state eGenes are pervasively depleted in GWAS for the 22 HCHF-related traits. Together, our results suggest that DR effects identified in baboons are translationally relevant to human health and can be leveraged to characterize the function of disease-associated genes.

## Discussion

In our two-year dietary intervention study, we observed considerable effects of a sustained HCHF diet on global gene expression levels in baboons and identified genetic loci that are associated with inter-individual transcriptional differences in response to diet. We found evidence for both sex- and tissue-specific transcriptional changes, reflecting the complex landscape of dynamic gene-by-diet interactions. We characterized hundreds of eQTLs in response to diet and found that diet-responsive eGenes in baboons can be valuable to interpret genetic associations with disease in humans. Our results further emphasize the intricate complexity of genetic regulatory effects and the value of considering GxE interactions in genetic studies of complex disease.

Our study underscores the strength of the baboon model in studying GxE interactions that present formidable challenges in human studies. The physiological similarity between baboons and humans, combined with the controlled environmental conditions of our study, enables us to suggest additional candidate genes associated with diseases that previously evaded identification in human studies of gene regulation.

For example, *APOA2* has a diet-specific genetic association with BMI and obesity in human populations^37–39^. Yet, despite having much larger sample sizes, no standard eQTLs for *APOA2* have been discovered in adipose, liver, or muscle tissues from GTEx^7^. Here, we identified an eQTL for the baboon *APOA2* ortholog in liver, which emerged solely in response to the HCHF diet. This example demonstrates the potential of contextually functional regulatory variants, even those found in non-human primates, in extending our understanding of disease loci. With the specific enrichment of DR eGenes among GWAS signals for human metabolic traits, the translational relevance of the baboon model to human health becomes undeniably clear.

By considering the genetic architecture of diet-responsive gene expression, we discovered that DR eQTLs exhibit a localization pattern that is distinct from non-DR eQTLs, featuring a more evenly distributed distance to the TSS. Disease-associated loci found in human GWAS also typically reside at greater distances from the nearest TSS compared to standard eQTLs^1^. Additionally, disease-associated loci in humans appear to evolve under stronger stabilizing selection^57,58^ and are associated with more complex regulatory landscapes than genes not associated with disease^6,66^. Similarly, compared to standard eQTLs, DR eQTLs are enriched among genes with complex regulatory landscapes, enduring stronger selective constraints, and participating in diverse biological functions. Together, our observations support the notion that context-specific eQTLs with more complex regulatory effects hold the key to a more profound understanding of GWAS signals.

It was previously shown that standard eQTLs generally evolve neutrally or under weak selection^67^, and that they are broadly shared across tissues^34^. Furthermore, genes that are highly conserved across species^63^, transcription factors, and genes in the center of regulatory networks^51^ are all notably depleted for standard eQTLs. It is therefore not surprising that our findings reveal systematic differences between context-responsive eQTLs and standard eQTLs. Not only do context-responsive eQTLs have more complex regulatory effects, but they are more likely to be functionally important and subject to natural selection, making them more likely to underlie inter-individual differences in disease risk.

We anticipate that expanding resources in non-human primates, even with relatively small available sample sizes, will enable further characterization of complex, multi-tissue disease-relevant GxE effects and enhance the interpretation of GWAS signals for a wide range of complex traits in humans. This, in turn, will provide additional support for the notion that eQTLs with disease-relevant GxE interactions are crucial for a better understanding of non-coding GWAS variants.

## Acknowledgments

We thank O. Allen for assistance with RNA extraction; J. O’Connell for advice on phasing; Y. Yang for help with ultimate deconvolution; B. Fair and B. Umans for feedback on the manuscript; N. Gonzales for editing and providing comments on the manuscript; C. Jones and S. Sumner for feedback on the general readability of the manuscript; University of Chicago Research Computing Center for providing computational resources. Figure 1A was created using BioRender. Y.G. and W.L. were supported by NIH/NIGMS R35 GM131726. L.C., J.V. and D.N. were supported by NIH/NHLBI P01 HL028972 and NIH P51 OD011133.

## Author contributions

W.L, Y.G., M.O., and L.C. conceptualized the study. L.C. and J.V. designed and conducted the animal work and diet study. D.N. cleaned and managed the animal physiological data. G.L. managed the tissue archive. W.L. performed the RNA extraction, RNA-seq experiments, analyzed the data, interpreted the results, and wrote the manuscript with contributions from Y.G and L.C. M.A. guided W.L. on genotype imputation. J.W. provided WGS data and guided W.L. on WGS data processing. Y.G. supervised and guided W.L.

## STAR★METHODS

### KEY RESOURCES TABLE

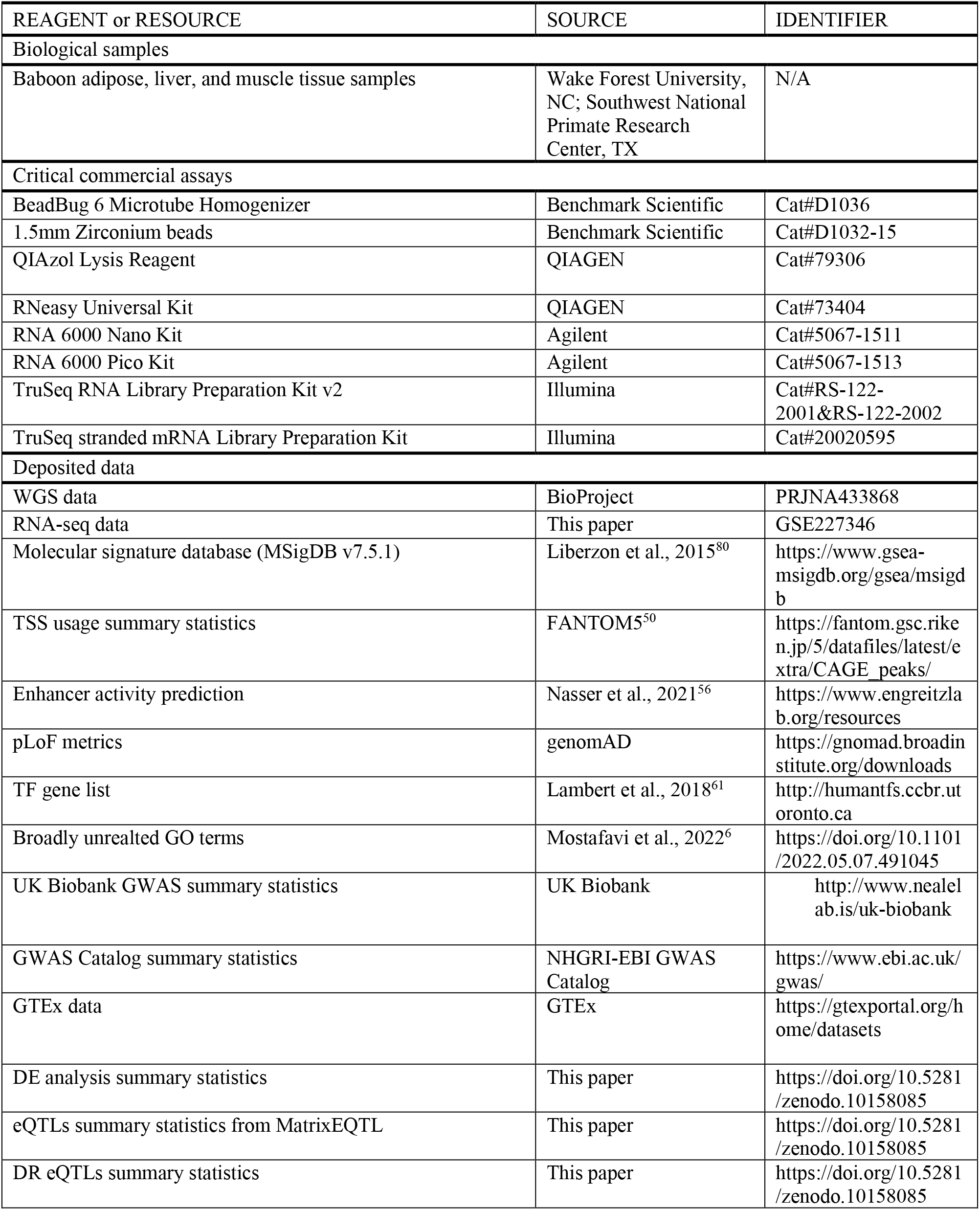
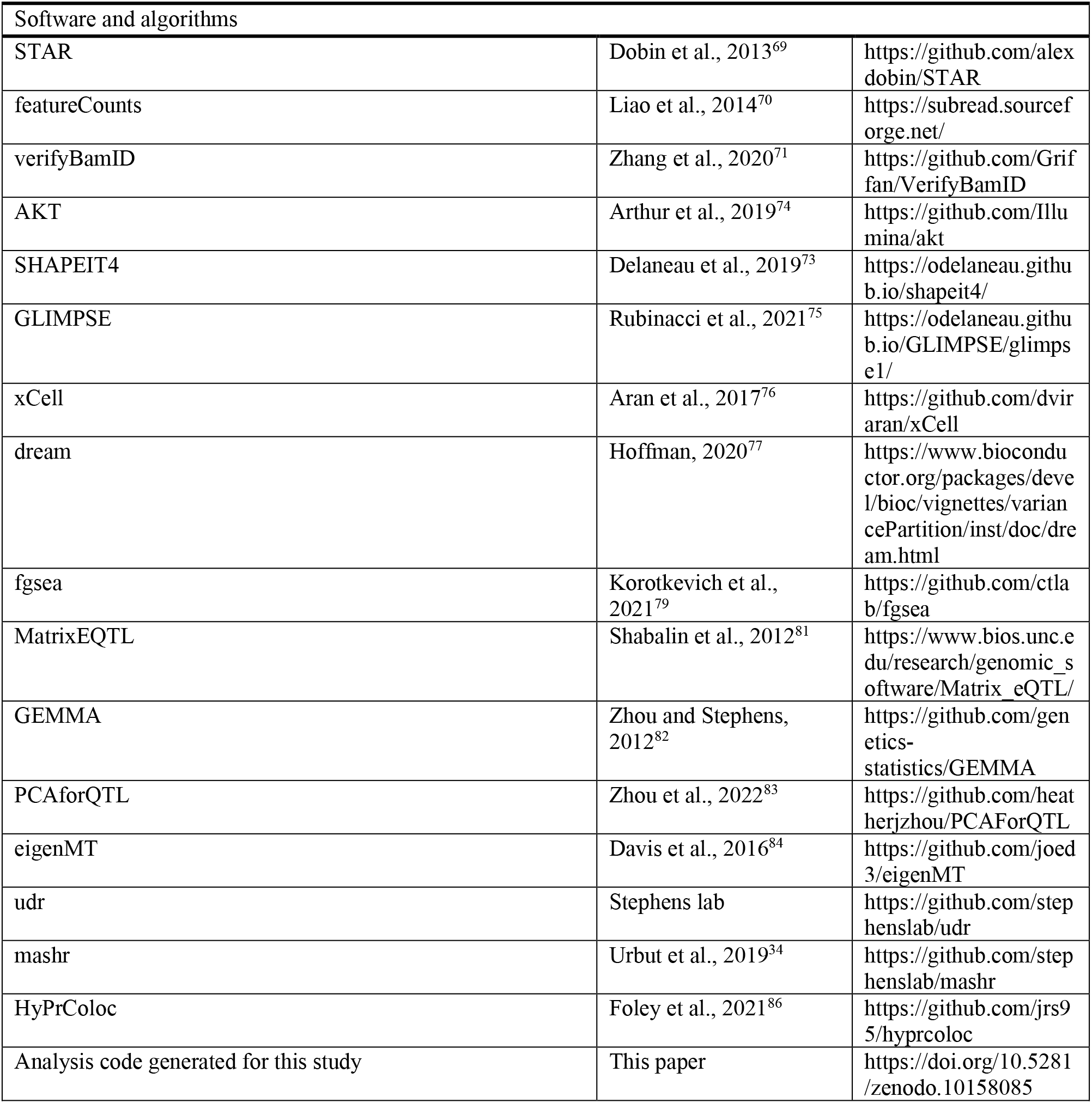

### RESROURCE AVAILABLITY

#### Lead contact

Further information and requests for resources and reagents should be directed to and will be fulfilled by the lead contact, Yoav Gilad.

#### Material availability

This study did not generate new unique reagents.

#### Data and code availability

The WGS data are available at BioProject. The RNA-seq data have been deposited at GEO. Accession numbers are listed in the key resources table. All original code, data, and summary statistics presented in this paper have been deposited at Zenodo. DOIs are listed in the key resources table. Any additional information required to reanalyze the data reported in this paper is available from the lead contact upon request.

### EXPERIMENTAL MODEL AND STUDY PARTICIPANT DETAILS

The subjects of the diet challenge were olive baboons (*Papio hamadryas anubis*), yellow baboons (*Papio hamadryas cynocephalus*), and their hybrid descendants, all of which were members of a large pedigreed breeding colony developed and maintained at the SNPRC. The 99 baboons included 42 females and 57 males with ages ranging from 6 years to 14.7 years. The mean age for the females was 10.4 years (range = 6.4 to 14.7 years) and that of the males was 9.2 years (range = 6 to 12.5 years). The female animals in this study were not pregnant or lactating during the two-year diet intervention. They did not receive any form of birth control and were living with infertile males in their normal social groups. The male baboons in this study were housed in social groups separate from the females and were not castrated. All animal procedures were reviewed and approved by the Texas Biomedical Research Institute’s (TBRI) Institutional Animal Care and Use Committee (IACUC). Southwest National Primate Research Center (SNPRC) facilities at TBRI and the animal use programs are accredited by Association for Assessment and Accreditation of Laboratory Animal Care International (AAALAC), operate according to all National Institutes of Health (NIH) and U.S. Department of Agriculture (USDA) guidelines, and directed by veterinarians (DVM). The SNPRC veterinarians made all animal care decisions. All animals were housed in group cages allowing them to live in their normal social groups with *ad libitum* access to food and water. Enrichment was provided on a daily basis by the SNPRC veterinary staff and behavioral staff in accordance with AAALAC, NIH, and USDA guidelines.

#### METHOD DETAILS

### Diet intervention

The baboons were maintained on a chow diet, low in cholesterol and fat (LCLF) from birth until they began a two-year dietary challenge with a diet high in cholesterol and saturated fat (HCHF). Table S1 shows the composition of the chow LCLF diet (Monkey Diet 15%/5LEO, LabDiet), the chow diet was used to prepare to atherogenic HCHF diet (Monkey Diet 25/50456, LabDiet), and the complete HCHF diet. To make the HCHF diet, we add a mix of lard, cholesterol, sodium chloride, vitamins [ascorbic acid and vitamin A (a retinyl acetate)], and water to the chow diet.

Metabolizable energy is approximately 3.8 kcal/g, with 40% of calories from fat, 40% of calories from carbohydrates, and 20% of calories from protein. We measured the composition of total fatty acids by gas-liquid chromatography of the fatty acid methyl esters [on DB-225 column (15 m), J&W Scientific]. Saturated fatty acids include myristic (1.7%), palmitic (24.9%), and stearic (17.9%); monounsaturated fatty acids include palmitoleic (2%) and oleic (38.7%); and polyunsaturated acids include linoleic (13.9%) and linolenic (0.9%). All baboons in the study were fed daily and allowed to eat *ad libitum*. We were not able to monitor and measure the amount of food consumed by each individual animal as they were maintained in social group housing, consistent with best practices for this species. The approximate mean daily intake per animal of the LCLF diet was 500 g (∼1500 kcal) and that for the HCHF diet was 400 g (∼1200 kcal). The mean amount of cholesterol consumed daily by animals on each of these diets was approximately 30 mg and 2230 mg, respectively.

#### Tissue sample collection

Baboon tissue samples were collected just prior to beginning the two-year HCHF diet challenge (while on the LCLF diet) and again at the end of the two-year HCHF diet challenge. The baboons were sedated with ketamine (10 mg/kg), given atropine (0.025 mg/kg), and intubated. Body weight was measured on a calibrated electronic balance and body length was measured between head and feet, with the animals lying on their back. Omental adipose biopsy samples were collected through the abdominal wall; liver biopsy samples were collected from the left lobe of the liver; and skeletal muscle biopsy samples were collected from the quadriceps. During post-biopsy recovery, analgesia was provided in the form of Stadol, 0.15 mg/kg, twice a day, for 3 days and ampicillin, 25 mg/day for 10 days.

#### Lipid biomarker measurement

We obtained data on circulating cardiovascular disease (CVD) risk factors, including lipids and lipoproteins, lipoprotein-related enzymes, and biomarkers of inflammation and oxidative stress from blood samples collected before and after the two-year HCHF diet intervention. We quantified the concentrations of the following serum lipids and lipoproteins. Total serum cholesterol (TSC) and triglyceride (TRIG) concentrations were determined enzymatically using commercial reagents in a clinical chemistry analyzer. High density lipoprotein cholesterol (HDLC) was measured in the supernatant after heparin Mn^+2^ precipitation, and non-HDLC (LDLC) was calculated as the difference between total and HDLC. Concentrations of apolipoproteins AI (APOAI), B (APOB), and E (APOE) were determined using an immunoturbidometric approach with commercial reagents in a clinical chemistry analyzer. We measured plasma oxidized LDL (oxLDL) concentrations (U/L) immunologically using a sandwich-style enzyme-linked immunoabsorbent assay (Mercodia Oxidized LDL ELISA; ALPCO Diagnostics). Lipoprotein size distributions were estimated as absorbance in large size particles minus absorbance in small size particles. ΔHDL (dHDL) was calculated as [(fractional absorbance, 8.6 to 10.5 nm)(fractional absorbance, 12 to 19 nm)]x1000; and ΔLDL (dLDL) was calculated as [(fractional absorbance, 27.2 to 28.4 nm)(fractional absorbance, 24.4 to 26 nm)]x1000^68^. Most samples were run at least twice, and the mean was used as the final measurement value.

#### RNA sample processing and sequencing

Approximately 10 mg of each frozen tissue sample was homogenized in 1ml QIAzol Lysis Reagent (79306, QIAGEN) using BeadBug 6 Microtube Homogenizer (D1036, Benchmark Scientific) with 1.5mm Zirconium beads (D1032-15, Benchmark Scientific). RNA was immediately extracted from the homogenate using QIAGEN RNeasy Universal Kit (73404, QIAGEN). RNA concentration and quality was measured using Agilent RNA 6000 Nano Kit and RNA 6000 Pico Kit (5067-1511 & 5067-1513, Agilent) on Agilent 2100 Bioanalyzer (Figure S1A). 96 libraries from 16 animals in batch 1 were performed using TruSeq RNA Library Preparation Kit v2 (RS-122-2001, RS-122-2002, Illumina) and were sequenced 50 base pairs, single-end using the Illumina HiSeq4000 according to manufacturer instructions with the goal of achieving at least 15 million reads per sample (Figure S1B). Each sequencing lane contained 24 multiplexed samples from different RNA extraction batches. 498 libraries from 83 animals in batch 2 were performed using TruSeq stranded mRNA Library Preparation Kit (20020595, Illumina) and were sequenced 50 base pairs, paired-end using the Illumina NovaSeq6000 according to manufacturer instructions with the goal of achieving at least 30 million reads per sample (Figure S1B). Each sequencing lane contained 84 multiplexed samples from different RNA extraction batches. Samples from the same animal were always included in the same RNA extraction, library preparation, and sequencing batches. Samples from different animals were randomized in every processing step to avoid potential batch effects.

#### RNA-seq data

RNA-seq data were aligned to the baboon reference genome Panubis1.0 with STAR v2.7.7a^69^. FASTQC was used to confirm that the reads were of high quality. Gene-level expression quantification was performed using featureCounts^70^. Quality controls were performed on two levels using a similar pipeline for GTEx^7^. On the sample level, we first identified and removed five sample swaps and contaminations where a sample was not matched to its genotype or contained a mixture of two or more samples using verifyBamID^71^. We removed 18 outlier samples based on expression profile using PCA. We checked sex correctness by examining gene expression for genes on the Y chromosome for each sample (Figure S2). On the gene level, gene expression values for all samples from a given tissue were normalized using the following procedure: 1) read counts were normalized between samples using trimmed mean of M-values (TMM); 2) genes were selected based on expression thresholds of ≥0.1 TPM in ≥10% of samples, and > 3 reads (unnormalized) in ≥10% of samples for both diet conditions; 3) expression values for each gene were inverse normal transformed across samples. After the quality controls, we obtained 570 high-quality RNA samples and 20,224 genes for downstream analyses.

#### Genotype and imputation

WGS data was collected and provided by the Wall lab, with most of the animals having a low coverage genome (Figures 2A)^72^. We generated a reference panel using SHAPEIT4 from 202 high-coverage (>10X) baboons from the same colony. SHAPEIT4 is a fast and accurate method for estimation of haplotypes (i.e. phasing) for SNP array and high coverage sequencing data^73^. To increase the accuracy of phasing, we incorporated pedigree information by including sets of pre-phased genotypes (i.e. haplotype scaffold) derived from pedigree data using akt^74^. We next imputed the animals with low-coverage WGS data (<10X) using GLIMPSE. GLIMPSE is a powerful tool to impute genotypes from low-coverage sequencing data^75^. We validated the accuracy of imputation by imputing a pseudo-low-coverage genome. Using 30X WGS data from a baboon that was not included in the reference panel, we downsampled the reads to 4X coverage. We imputed this downsampled data and computed the squared Pearson correlation between imputed dosages (in MAF bins) and highly-confident genotype calls from the original high-coverage data (Figure S3B). For SNPs with MAF>5%, the correlation is larger than 0.9. Due to the lack of genetic maps for X and Y chromosomes, we were not able to phase or impute genotypes for the sex chromosomes, thereby including only autosomes. After excluding low allele-frequency (MAF < 5%) and monomorphic SNPs, we obtained 16,671,556 autosomal SNPs across 99 animals for downstream analyses.

#### Cell type enrichment

Cell type enrichment scores for each sample were computed by running xCell on the full TPM gene expression matrix of the 570 post-QC samples. The xCell method uses single-sample geneset enrichment analysis (ssGSEA) to score each sample and estimate the enrichment of 64 reference cell types, spanning multiple adaptive and innate immunity cells, hematopoietic progenitors, epithelial cells, and extracellular matrix cells. The reference cell types are defined by gene signatures learned from thousands of pure cell types from various sources^76^. The raw enrichment score per cell type is defined as the average ssGSEA score from all the cell types’ corresponding signatures. Then, the raw scores are transformed to a linear scale that resembles percentages. The cell type enrichment scores shown in the results are transformed scores. For sanity check, we examined the enrichment of representative cell types for each tissue including adipocytes, hepatocytes, and myocytes (Figure 1B). For each tissue, we fit a linear regression to test for differences in the cell type enrichment between diets while adjusting for sex, age, RIN, and batch effects.

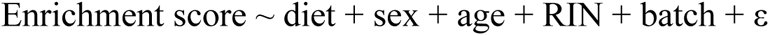

We corrected for multiple testing using the Benjamini-Hochberg procedure and determined significance as FDR ≤ 0.05. We visualized sample clustering using heatmaps for the 15 highly enriched cell types with both mean and median enrichment scores > 0.05 in at least one tissue (Figure S7A). We calculated correlations between the enrichment and age, sex, and diet in each tissue using Spearman’s correlation test (Figure S7B).

#### Differential expression

We performed differential expression (DE) analysis using dream. Dream uses a linear mixed model to increase power and decrease false positives for RNA-seq datasets with repeated measurements^77^. In each tissue type, we set diet, sex, age, and RIN as fixed effects and individual and batch as random effects in a linear mixed model. We first used sva^78^ to identify surrogate variables for unknown sources of variation. Using the function num.sv, we identified zero latent factors that need to be estimated and accounted for. Using dream, we estimated precision weights, fit the model on each gene using the Satterthwaite approximation for the hypothesis test by default, and applied empirical Bayes shrinkage on the linear mixed model.

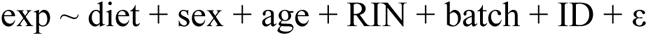

The response variable ‘exp’ is a vector of TMM-normalized expression level of all samples for a given gene. ε is normally distributed error modeled with precision weights via an iterative optimization algorithm. Hypothesis testing is performed by specifying a contrast matrix that is a linear combination of the estimated coefficients and evaluating the null model. Genes with an FDR-adjusted p value < 0.01 were considered DE genes. We applied mash to estimate sharing of DE across tissues. mash is a powerful statistical method to estimate and test for effects across conditions while accounting for the sharing information^34^. For all tested genes, we provided the effect size and corresponding standard error matrices estimated from dream to mash to jointly model differential expression. We filled missing values with an effect size of 0 and a standard error of 10 for genes excluded for low expression in a particular tissue. We estimated paired-wise sharing by LFSR < 0.01 in at least one condition and the magnitude of effect sizes > 1.5 between conditions from mash (Figure S4B).

#### Sex-biased differential expression

We performed sex-biased differential expression analysis using the same procedures. We defined sex-biased diet-responsive (DR) effect on gene expression as difference in diet effect on gene expression between males and females. Specifically, when the expression level of a gene is altered in response to diet, and when this altered response differs between males and females, we call this a sex-biased diet-responsive effect on gene expression. We fit a linear mixed model using dream while accounting for age, RIN, batch and individual effects.

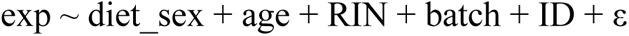

In each tissue type, we set diet_sex (combination of diet and sex), age, RIN, and batch as fixed effects and individual as a random effect in a linear mixed model. dream allows us to perform a hypothesis test of a linear combination of coefficients by using a contrast matrix from the ‘makeContrastsDream’ function in R package variancePartition. Here, we tested the difference of diet effect between males and females (i.e. sex-biased diet effects), asking which genes respond differently to the HCHF diet in males compared to females. The difference of differences is the interaction term in the model. Due to the smaller effective sample size in sex-biased DE analysis, we used FDR < 0.05 to identify sex-biased DE genes.

We replicated the results by using another approach where we searched for sex-specific DE genes in each tissue (Figure S8). We performed DE analysis for males and females separately in each tissue using the same model as described in the “differential expression” section above. We performed joint analysis by feeding the statistics from the sex-stratified DE analysis to mashr where we treated each sex and tissue combination as conditions. We identified sex-specific DE genes in each tissue by local false sign rate (LFSR) < 0.05 in at least one condition and the magnitude of effect sizes > 2.5 between diets.

#### Gene set enrichment analysis

We performed gene set enrichment analysis (GSEA) using the R package fgsea. fgsea is a powerful method that quickly estimates arbitrarily low GSEA p-values accurately based on an adaptive multi-level split Monte-Carlo scheme^79^. We excluded unannotated genes with a symbol beginning with “LOC” whose human orthologs have not been determined in the baboon reference genome (Panubis1.0). We used pre-ranked t-statistics for all tested annotated genes in each tissue from DE analysis as input and 50 hallmark gene sets from the Molecular Signature Database (MSigDB v7.5.1) as reference gene sets^80^. The hallmark gene sets summarize and represent specific well-defined biological states or processes and display coherent expression. These gene sets were generated by a computational methodology based on identifying overlaps between gene sets in other MSigDB collections and retaining genes whose gene expression levels are coordinate. Multiple testing was corrected using the Benjamini-Hochberg procedure and an FDR ≤ 0.05 and absolute NES > 1 was considered as significant enrichment. NES is a normalized enrichment score used to account for the size of a gene set and correlation between a gene set and genes in the ranked list.

#### eQTL mapping

cis-eQTL mapping was performed using a relatedness-accounted modified linear regression model implemented by MatrixEQTL^81^. In a regular linear regression model, the error term ε is assumed to be independent and identically distributed (i.i.d) across samples (i.e. uncorrelated homoscedastic errors).

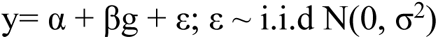

To account for the relatedness of the samples, we used correlated errors by feeding a relatedness matrix (K) as the error variance-covariance matrix to the parameter *errorCovariance* in MatrixEQTL.

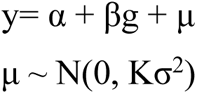

To maintain the assumption of a linear regression model, MatrixEQTL makes an internal transformation on the input variables, which makes the errors independent and identically distributed.

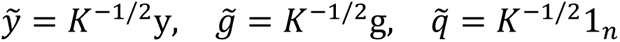

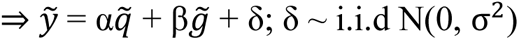

The genetic relatedness matrix was estimated from all 99 animals’ genotypes using GEMMA^82^ and it reflects the genetic relatedness relationship between each pair of individuals. We validated the relatedness estimation by comparing with pedigree information. The matrix was subset by the animals included in each condition and if necessary, the matrix was transformed to positive definite using the *make.positive.definite* function from the R package corpcor for the eQTL mapping.

We filtered SNPs in each tissue and diet combination by allele frequency and genotype frequency. We removed outlier SNPs with MAF < 0.05. We also filtered out loci with < 2 homozygotes across samples in each tissue and diet combination.

We included the known variables sex, age, RIN, and batch as covariates. To capture hidden technical or biological factors of expression variability, we infer principal components (PCs) from the gene expression matrix for each tissue x diet condition using PCAforQTL^83^. The number of PCs was selected and included in the model as covariates based on the elbow method and the Buja and Eyuboglu (BE) algorithm, which also approximately maximized cis-eGene discovery. We computed the correlation of the selected PCs and the known variables and removed those that were highly correlated with any of the selected PCs (R^2^ > 0.9).

The mapping window was defined as 100kb up- and down-stream of the transcription start site. We output all results by setting the *pvOutputThreshold*=1. To identify genes with at least one significant eQTL (cis-eGenes), the top nominal p-value for each gene was selected and corrected for multiple testing at the gene level using eigenMT^84^. eigenMT is an efficient multiple hypothesis testing correction method that estimates the effective number of tests using a genotype correlation matrix and then applying Bonferroni correction. Genome-wide significance was then determined by computing Benjamini-Hochberg FDR on the top eigenMT-corrected p-value for each gene.

#### Diet-responsive eQTLs

To combat the issue of incomplete power, we used mash to estimate sharing between conditions, and to identify diet-specific dynamic effects in each tissue. By jointly analyzing genetic effects across multiple conditions, mash increases power and improves effect-size estimates, thereby allowing for greater confidence in effect sharing and estimates of condition-specificity. We ran mashr using the output of eQTL mapping from MatrixEQTL of all six conditions. To avoid potential bias of including tissue-specific genes in the joint analysis for discovering tissue-specific effects, we included commonly shared genes across all conditions. No effect size is *post hoc* set to 0 for mashr. To better learn the heterogenous patterns of sharing from data and improve the accuracy of estimates, we implemented an optimized algorithm, ultimate deconvolution^85^. For each gene, the top SNP with the largest univariate |Z|-statistic across all conditions was selected. The top SNPs with LFSR < 0.05 were designated as significant eQTLs. To investigate sharing of the top eQTLs between diets, we assessed sharing of effects by magnitude (effects have similar magnitude within a factor of 1.5) for the top SNPs that are significant eQTLs in at least one diet within each tissue separately. DR eQTLs are SNPs that are significant eQTLs in at least one diet for a given tissue and the fold change of effects between diets is more than a factor of 1.5. A DR eGene is a gene with at least one DR eQTL. Non-DR eQTLs are SNPs that are significant eQTLs in at least one diet for a given tissue and the fold change of effects between diets is less than a factor of 1.5. A non-DR eGene is a gene with at least one eQTL but no DR eQTLs.

#### Physiological QTL mapping

We removed outliers by dropping any sample that had a biomarker level greater than 20-fold the interquartile range or greater than 10-fold below the median across all samples. The measurement values for each biomarker were inverse normal transformed across samples. We mapped physiological quantitative trait loci using the same genotype data and model as eQTL mapping pipeline described above, this time testing the entire genome rather than just the cis eQTL region.

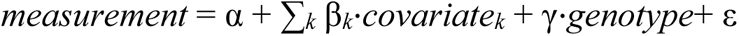

#### Colocalization

We used HyPrColoc to determine whether eQTL and trait associations are explained by the same causal variant explains. HyPrColoc is an efficient deterministic Bayesian divisive clustering algorithm that uses GWAS summary statistics to detect colocalization across vast numbers of traits simultaneously^86^. We used the default setting of prior probability = 1×10^-4^ and a conditional colocalization prior = 0.025. We defined a significant colocalization as having a posterior probability larger than 0.5.

#### Sex-biased eQTLs

We define sex-biased eQTLs as those cis-eQTLs with a significantly different genetic effect between males and females. We modified the relatedness-accounted linear regression model described in the eQTL mapping section by adding a genotype-by-sex (G x Sex) interaction term and tested for significance of G x Sex interaction. This was implemented in MatrixEQTL by setting *useModel = modelLINEAR_CROSS.* We filtered SNPs in each tissue and diet combination by removing outlier SNPs with MAF < 0.05 and loci with < 2 homozygotes across samples in each sex group. We used the same covariates, mapping window, and multiple testing correction approach as described above.

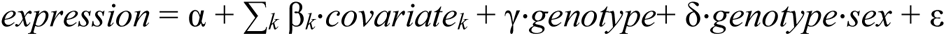

#### Comparison of genic features

We compared genic features of DR eGenes, nonDR eGenes, and all eGenes in each tissue. We calculated the TSS count of a given gene by the number of promoter peaks measured by the FANTOM5 project using Cap Analysis of Gene Expression (CAGE). We estimated enhancer activity of a given gene by computing the accumulative enhancer length across 131 cell or tissue types predicted based on the activity-by-contact (ABC) model (ABC score > 0.05). We defined loss-of-function (LoF)-intolerant genes by pLI score > 0.9 using the statistic from gnomAD.v2.1.1. For Gene Ontology (GO) analysis, we focused on 41 broadly unrelated GO terms that are used in Mostafavi et al. For enrichment in transcription factors (TFs), we used the official list of human TFs (v1.01) from Lambert et al. which includes 1639 human TFs. For all the analyses, we only included the genes that are shared between the testing set for eQTL mapping in this study (n=14814) and the reference set of the interest. Enrichment tests were performed using Fisher’s exact test.

#### Enrichment with human GWAS

We obtained GWAS summary statistics from UK Biobank provided by Ben Neale’s lab^64^ and The NHGRI-EBI Catalog of human GWAS (v1.0.2, 2022-10-08)^65^. For GWAS from UK Biobank, we selected 22 HCHF-related traits or disease and linked each GWAS hit to the closest TSS among 19393 protein-coding genes from GENCODE V43. For GWAS from GWAS Catalog, we obtained 507 traits using the “MAPPED_TRAIT” column after filtering for traits with at least 30 associated genes. We classified the traits into metabolic-relevant (n=98) and non-metabolic relevant traits (n=213). Ambiguous traits for the classification such as immune- and blood-relevant traits were removed. We obtained genes associated with each trait using the “MAPPED GENE(S)” column where if a SNP is located within a gene, that gene is listed; if a SNP is located within multiple genes, all genes are included; if a SNP is intergenic, the closest gene is included. We tested for enrichment of baboon DR eGenes in human GWAS genes for each trait using Fisher’s exact test. We used Fisher’s method to calculate the combined p-values for metabolic-relevant traits and non-metabolic relevant traits, respectively.

