## Supplementary material for "Genetic regulatory effects in response to a high cholesterol, high fat diet in baboons": Figs. S1-S17, Table S1

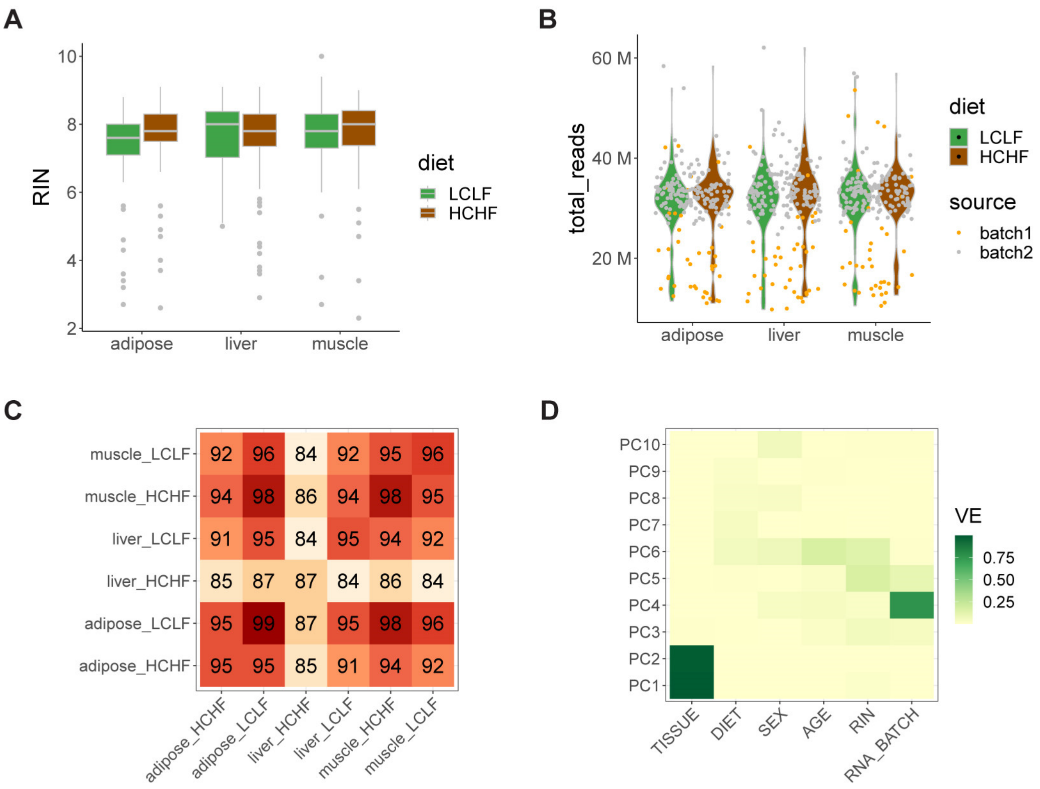


**Figure S1. Overview of the RNA samples.**

(**A**) RNA quality measured by RNA integrity number (RIN) of 594 RNA samples.
(**B**) Library size of RNA-seq samples in million of reads.
(**C**) Sample overlap after quality control (n=570).
(**D**) Variance of the first ten principal components explained by known variables.

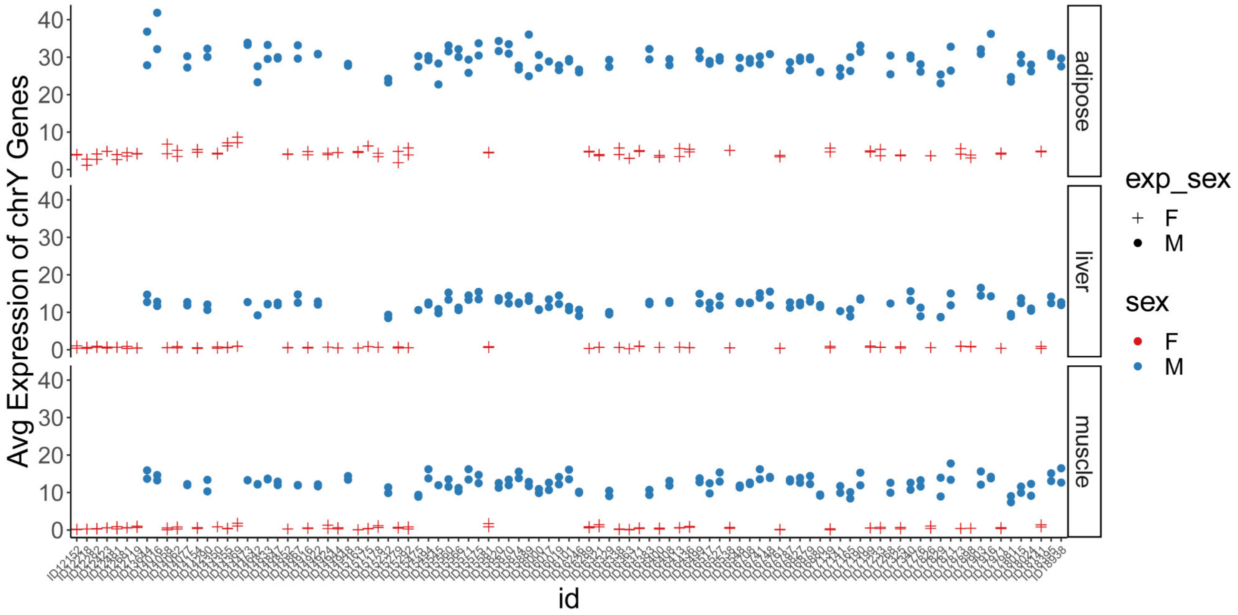


**Figure S2. Sex confirmation by expression level of chromosome Y genes.**

Average expression level of all chromosome Y genes for each sample. Sex inferred by gene expression is indicated by different shapes and the sex information from animal metadata is denoted by color.

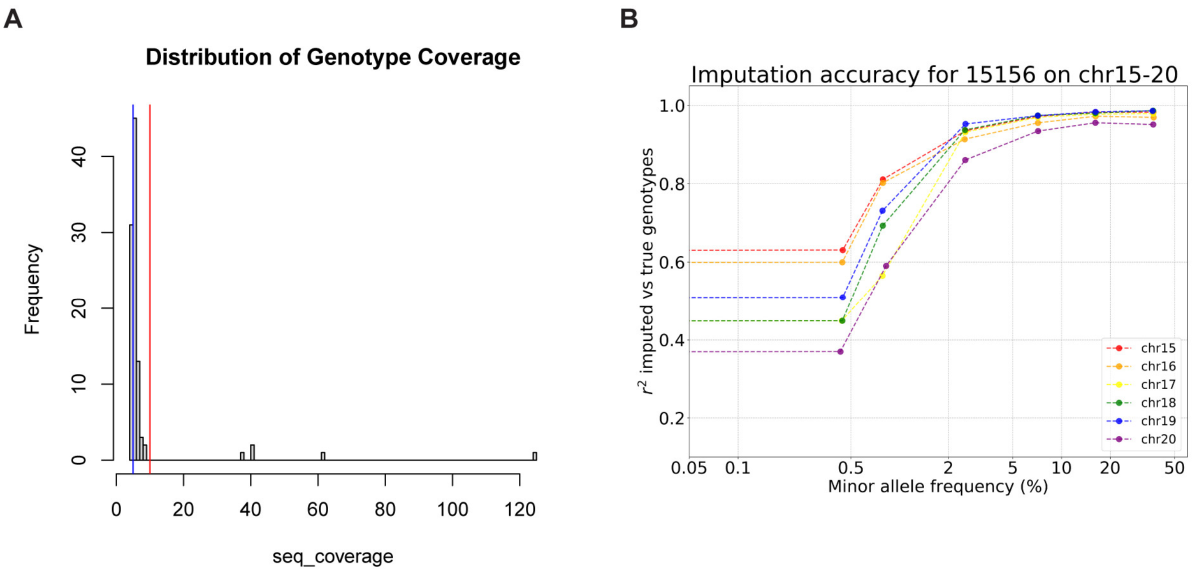


### Figure S3. Whole-genome sequencing (WGS) data.

(**A**) Distribution of sequencing coverage of the WGS data for all 112 animals. The blue vertical line depicts 5X, and the red vertical line depicts 10X.

(**B**) Accuracy of low-coverage sequencing imputation using GLIMPSE. Genotype concordance to 30X coverage for the 4X coverage was estimated by aggregated $r^2$ stratified by minor allele frequency (MAF>0.5%) on chromosome 15-20.

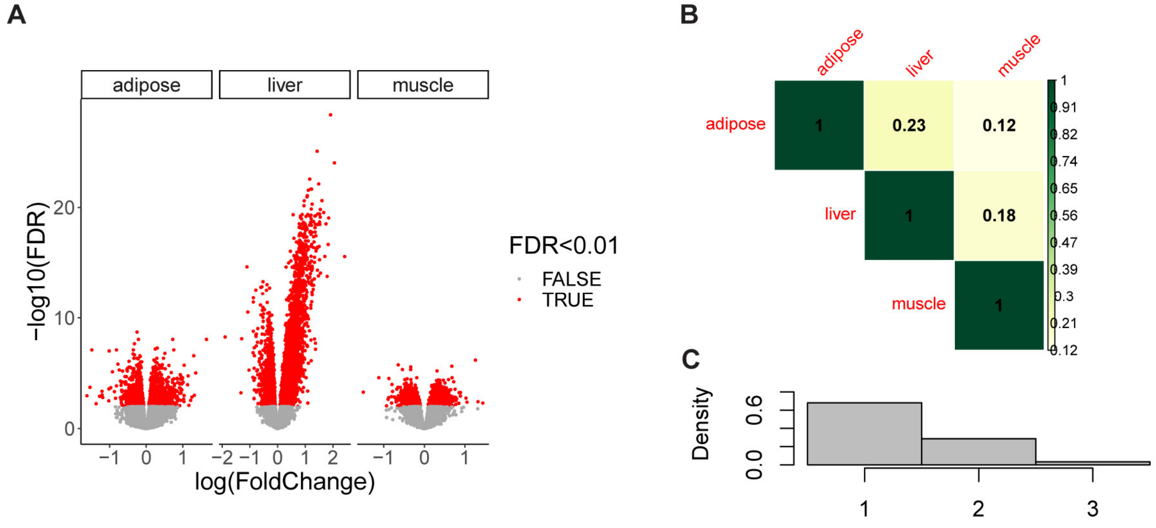


**Figure S4. Differential expression analysis in response to HCHF diet.**

(**A**) Volcano plot for DR differential expression. Red dots are significant DR genes with an FDR threshold of 0.01.
(**B**) Sharing of DR genes between tissue after joint analysis using mash.
(**C**) Distribution of the number of tissues where DR genes are shared.

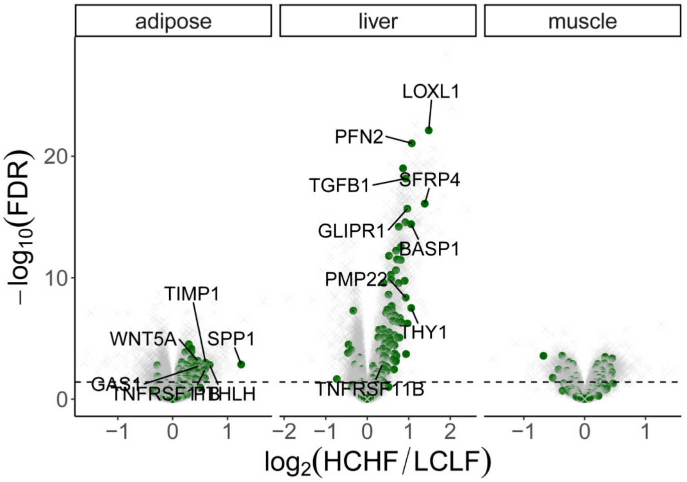


**Figure S5. Diet-responsive differential expression highlighted by genes involved in EMT (green).** DR genes are significantly enriched in EMT in adipose ($P < 1 \times 10^{-16}$) and liver ($P < 1 \times 10^{-15}$), but not in muscle ($P = 0.6$). The dashed line indicates an FDR threshold of 0.01.

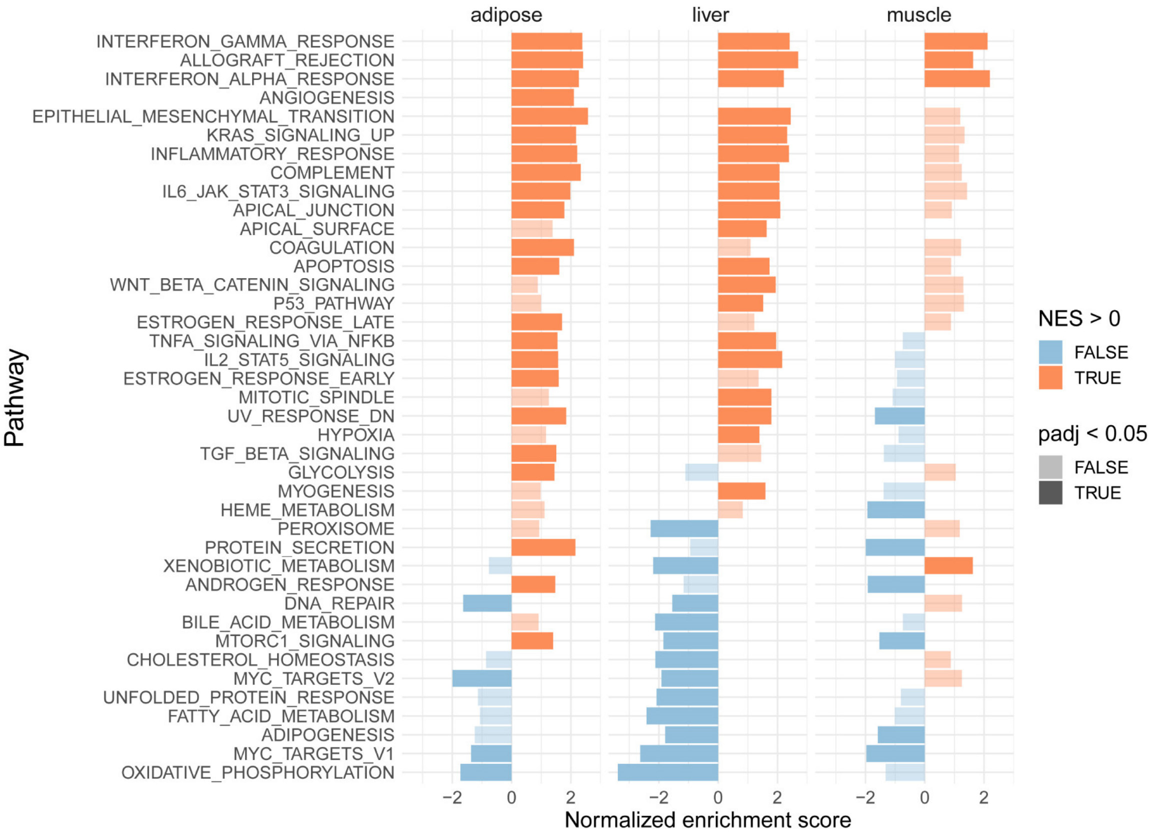


**Figure S6. Pathway enrichment of diet-responsive genes in adipose, liver, and muscle.** The colors denote direction of enrichments and opacity of the color indicates the statistical significance. For opaque colors, the normalized enrichment score (NES) FDR ≤ 0.05 and for transparent colors, the NES FDR > 0.05.

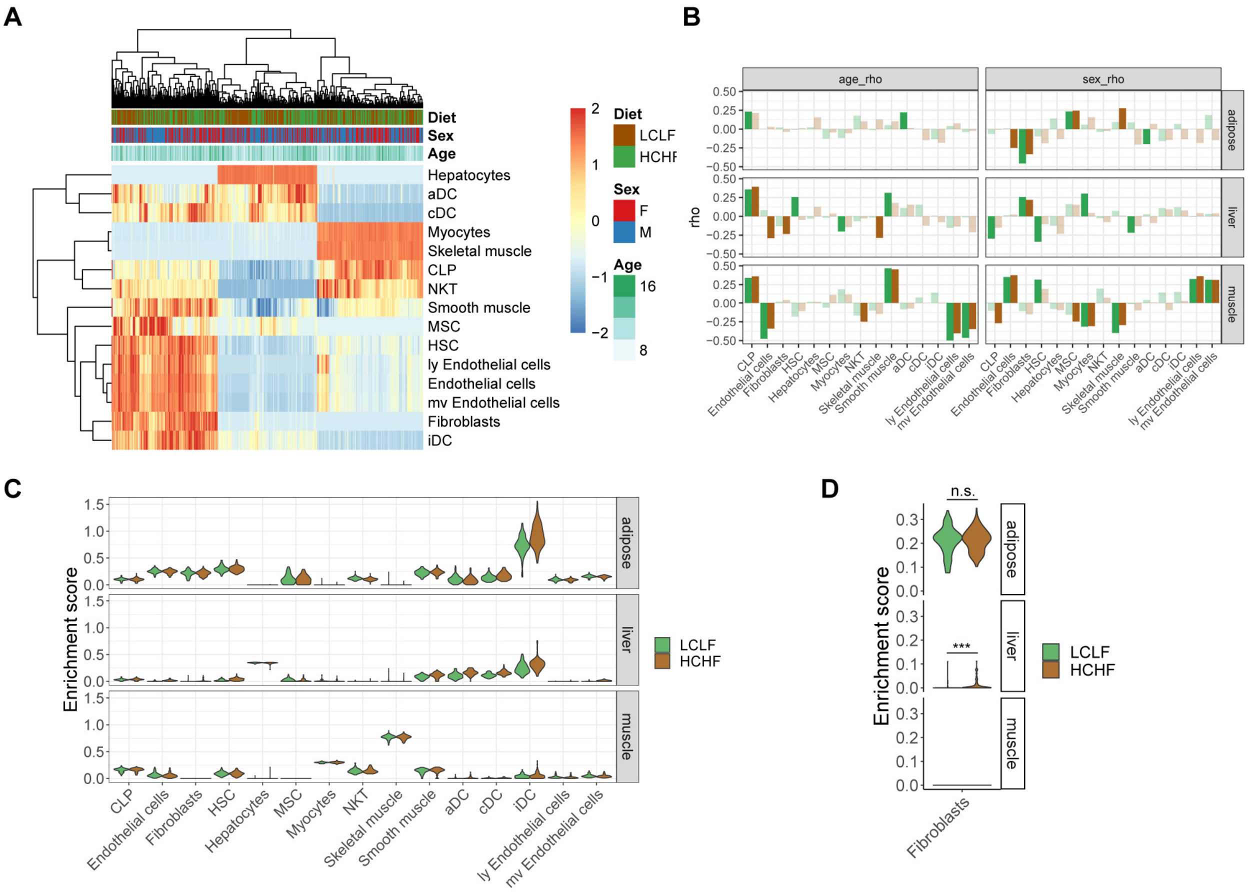


**Figure S7. Cell type enrichment analysis in response to HCHF diet.**

(**A**) Heatmap of samples by the inferred enrichment scores for the primary cell types.
(**B**) Spearman's correlation (rho) of the inferred enrichment scores for the primary cell types with age or sex, grouped by the diet condition. Opaque colors are significant at an FDR threshold of 0.05; transparent colors are not significant at an FDR threshold of 0.05.
(**C**) Comparison of cell type enrichment between diet conditions in abundant cell types (total enrichment score >15 in at least one tissue type) , including common lymphoid progenitors (CLP), endothelial cells, fibroblasts, hematopoietic stem cells (HSC), hepatocytes, mesenchymal stem cells (MSC), myocytes, natural killer T-cells (NKT), skeletal muscle cells, smooth muscle cells, activated dendritic cells (aDC), conventional dendritic cells (cDC), immature dendritic cells (iDC), lymphatic (ly) endothelial cells, and microvascular (mv) endothelial cells.
(**D**) Fibroblast enrichment changes in response to the HCHF diet. Asterisks indicate statistical significance (ns: non-significance, ***P < 0.001).

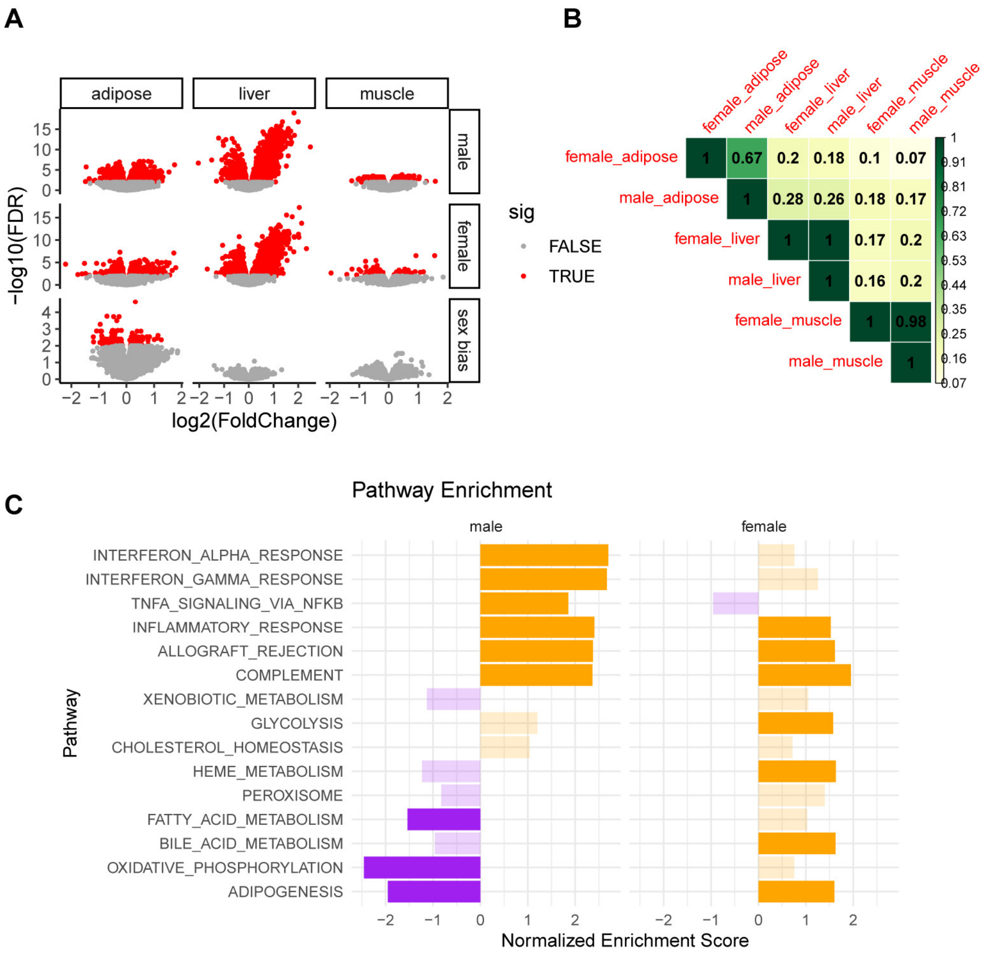


**Figure S8. Sex-stratified DE analysis in response to diet.**

(**A**) DE analysis of diet response in males (top), diet response in females (middle), and sex-biased diet response (bottom). Red dots are significant DR genes with an FDR threshold of 0.01.
(**B**) Pairwise sharing of male DR genes and female DR genes across sex and tissue combinations after joint analysis.
(**C**) Pathway enrichment of DR genes in males and females, separately. The colors denote direction of enrichments and opacity of the color indicates the statistical significance. Opaque colors are significant at an FDR threshold of 0.05; transparent colors are not significant at an FDR threshold of 0.05.

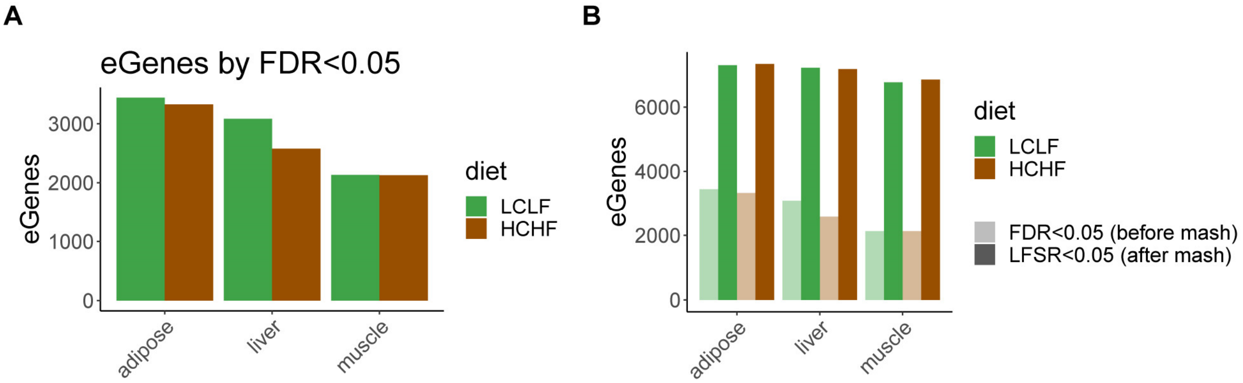


**Figure S9. Standard eGenes discovery in each diet and tissue combination.**

(**A**) The number of eGenes identified from MatrixEQTL with an FDR threshold of 0.05.
(**B**) Comparison of the number of eGenes identified in each diet condition using a linear model (FDR < 0.05) and after joint analysis using mash (LFSR < 0.05). eGenes identified using a linear model are shown in light colors and eGenes identified by mash are shown in dark colors.

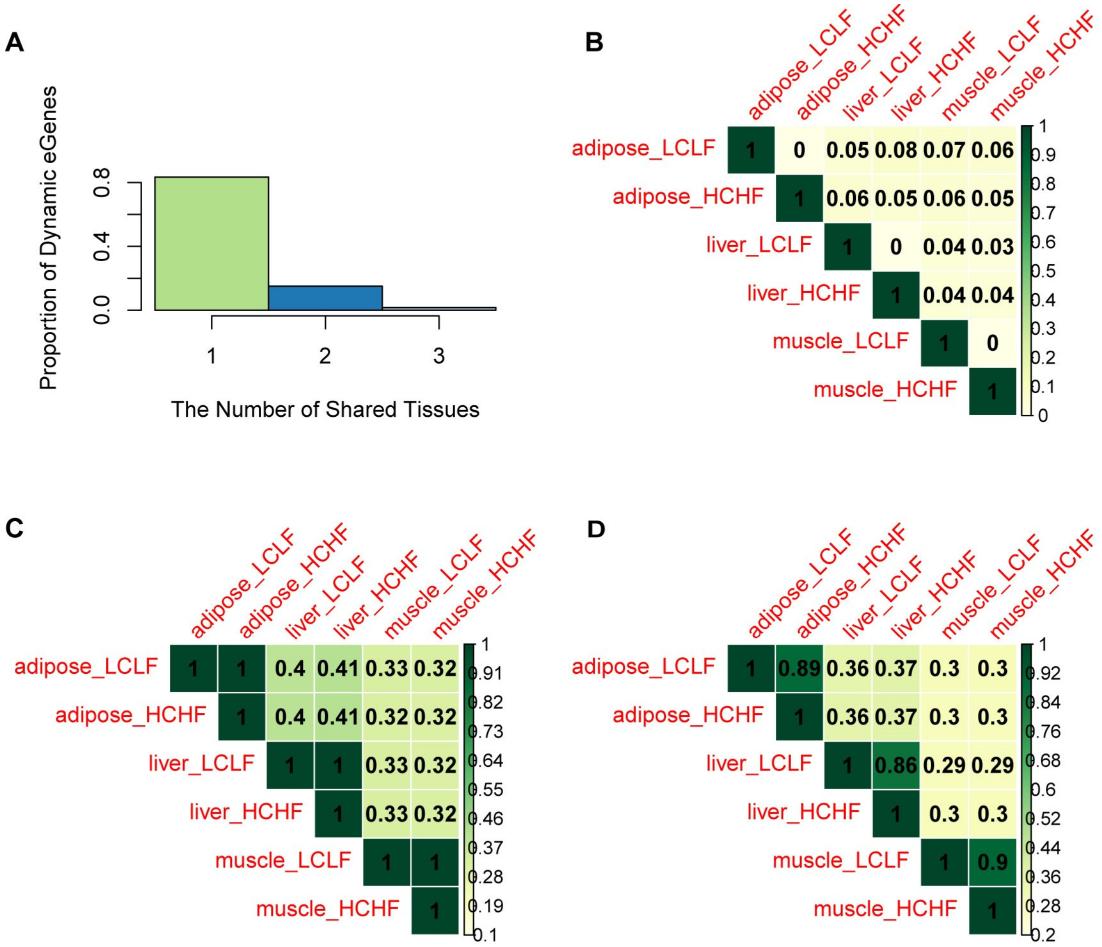


**Figure S10. Tissue-sharing of DR eQTLs and steady-state eQTLs.**

(**A**) Distribution of the number of tissues where DR eQTLs are shared.
(**B**) Pairwise sharing of DR eQTLs across diet and tissue combinations.
(**C**) Pairwise sharing of steady-state non-DR eQTLs across diet and tissue combinations.
(**D**) Pairwise sharing of all eQTLs across diet and tissue combinations

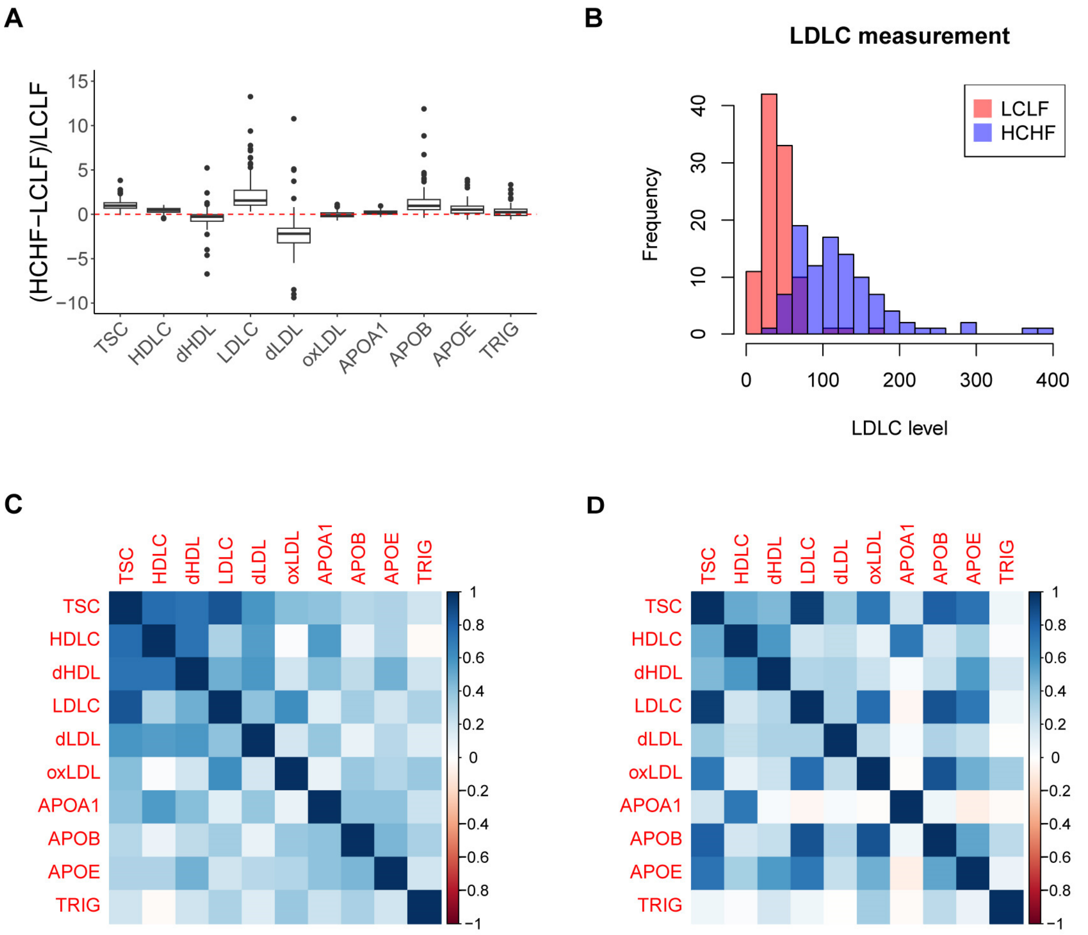


#### Figure S11. Overview of lipid biomarkers.

(**A**) Fold change of the measurement level of the 10 lipid biomarkers: total serum cholesterol (TSC), high density lipoprotein cholesterol (HDLC), low density lipoprotein cholesterol (LDLC), size distributions of HDL cholesterol (dHDL) and LDL cholesterol (dLDL), oxidized LDL (oxLDL), triglycerides (TRIG), and protein levels of APOA1, APOB, and APOE.
(**B**) Distribution of LDLC level, colored by diet condition.
(**C**) Correlation of the measurement level of the 10 lipid biomarkers in LCLF.
(**D**) Correlation of the measurement level of the 10 lipid biomarkers in HCHF.

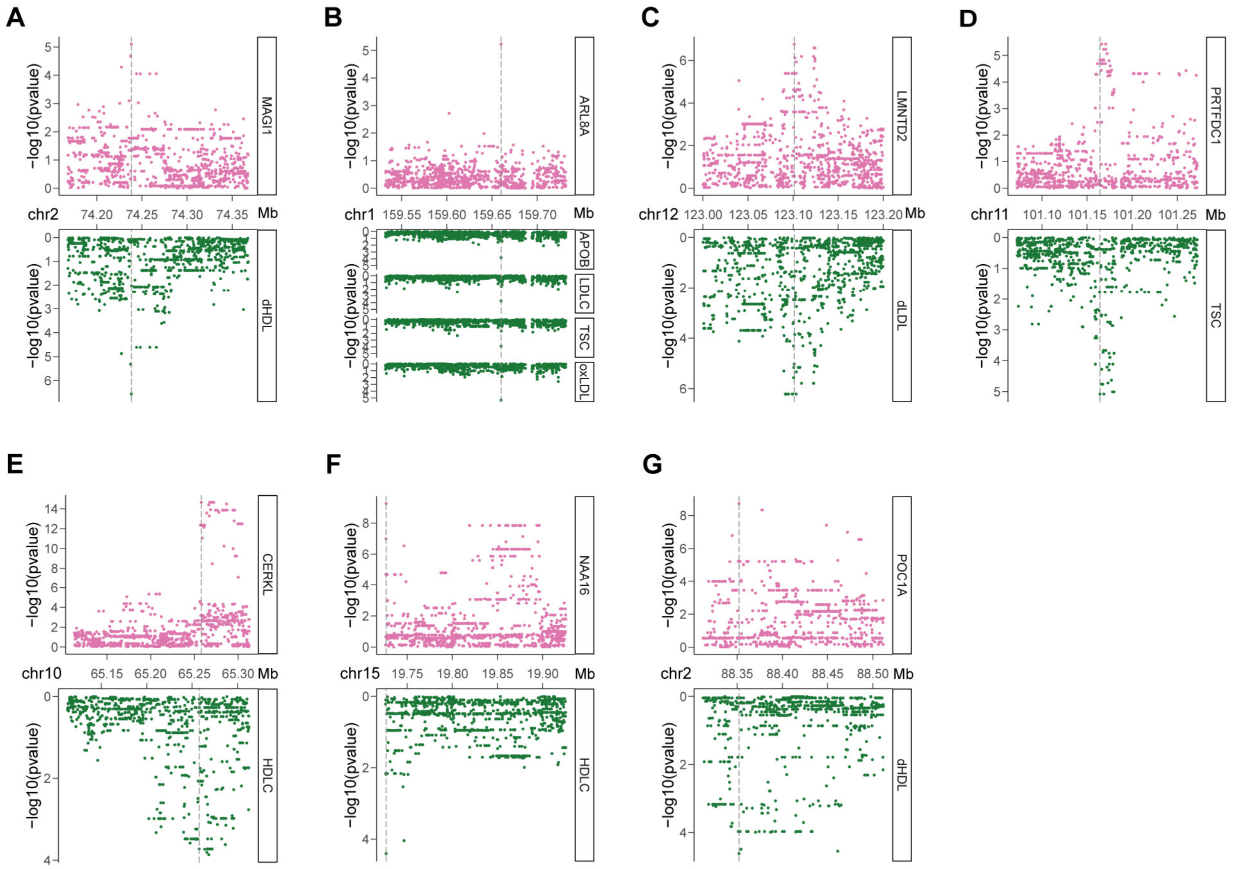


### Figure S12. Colocalizations of eQTLs and physiological lipid traits.

Visualization of all seven eQTL-lipid trait colocalizations (PP>0.5) where the top panel (pink) shows significance levels of variants evaluated as eQTLs for the given gene expression including all variants within 100kb of the transcription start site, and the bottom panel (green) shows significance levels of variants tested for association with the trait(s) within the same region. Vertical lines depict the genomic location of the candidate colocalized variant.

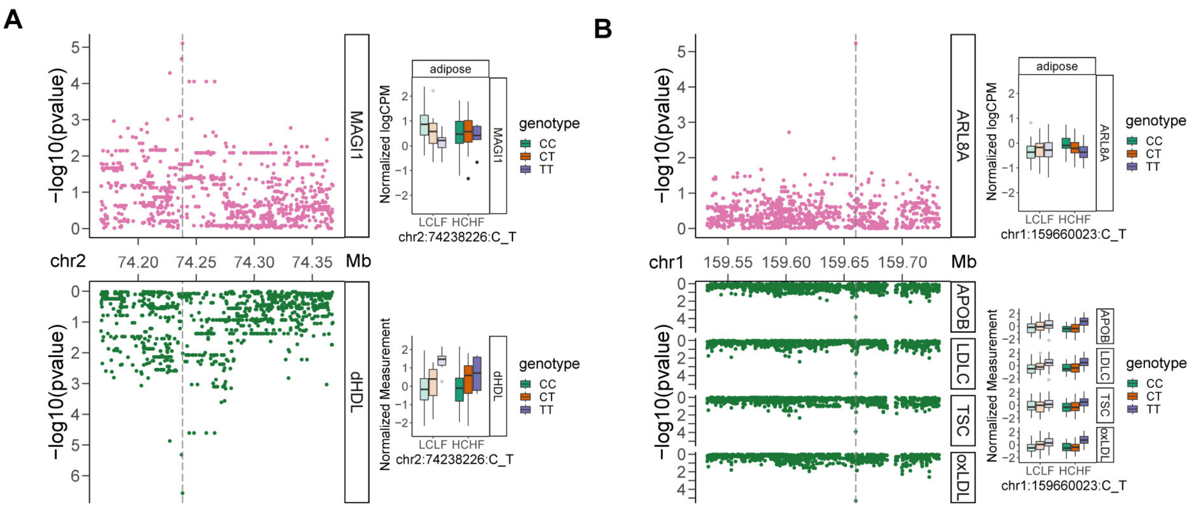


**Figure S13. DR eQTL-lipid trait colocalization.**

Colocalization of (**A**) DR eQTL for *MAGI1* with dHDL; (**B**) DR eQTL for *ARL8A* with APOB, LDLC, TSC, and oxLDL. Manhattan plots (left) show significance levels of variants tested for association with gene expression level (top) or with physiological lipid traits(s) (bottom). Vertical lines depict the genomic location of the candidate colocalized variant. Box plots (right) show association between genotype (color) of the colocalized locus and diet conditions on normalized gene expression (top) or normalized level of the lipid biomarker(s) (bottom).

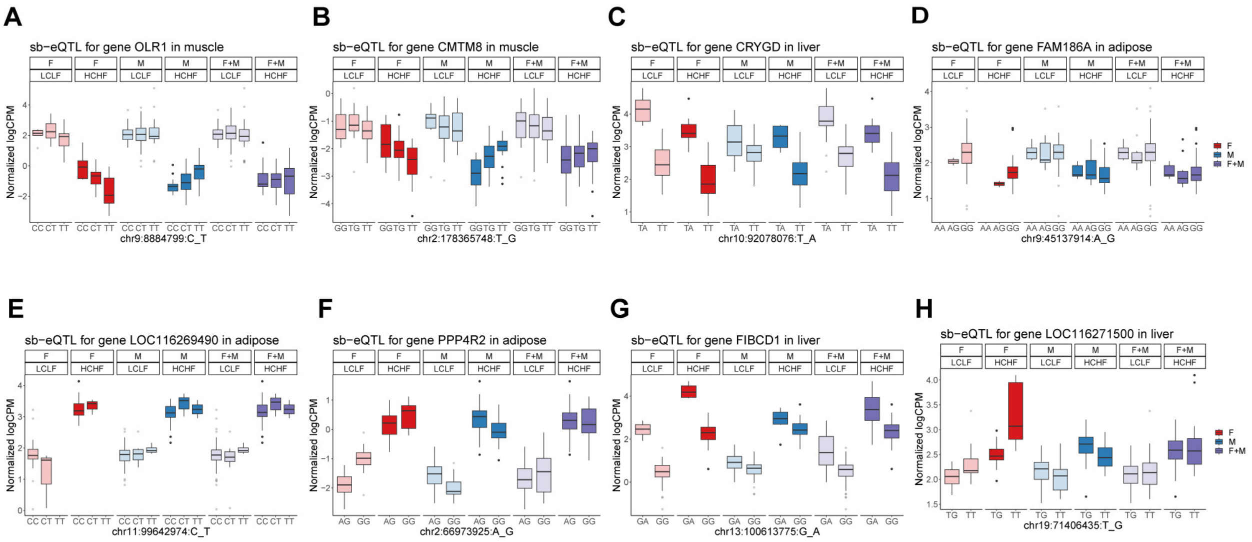


### Figure S14. Sex-biased eQTLs.

Visualization of all eight sex-biased eQTLs (FDR < 0.25) by diet condition (opacity) and sex group (color). (**A**)-(**E**) are sex-biased eQTLs that emerge only in response to one diet.

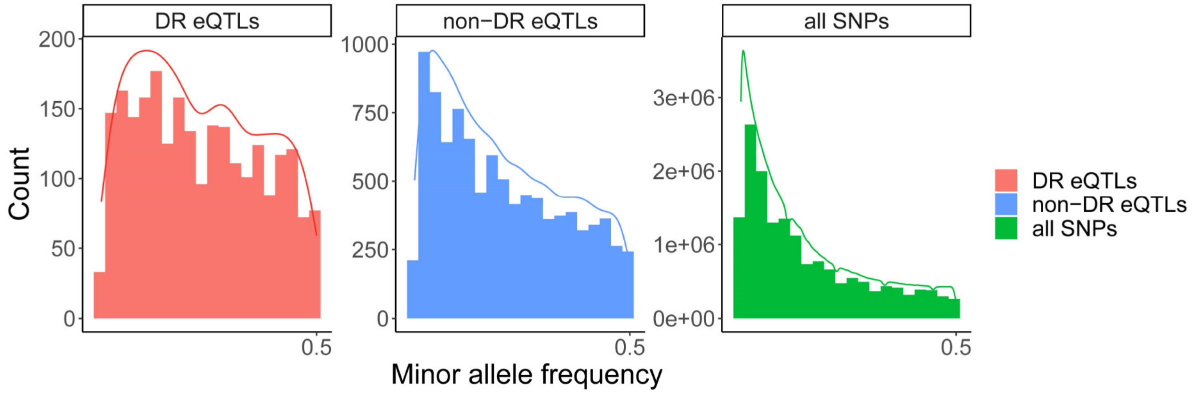


**Figure S15. Distribution of minor allele frequency (MAF).**

MAF of DR eQTLs (red), non-DR eQTLs (blue), and all SNPs (green). The histograms show MAF distribution in each group of SNPs. The curves show the kernel density estimate for each group of SNPs.

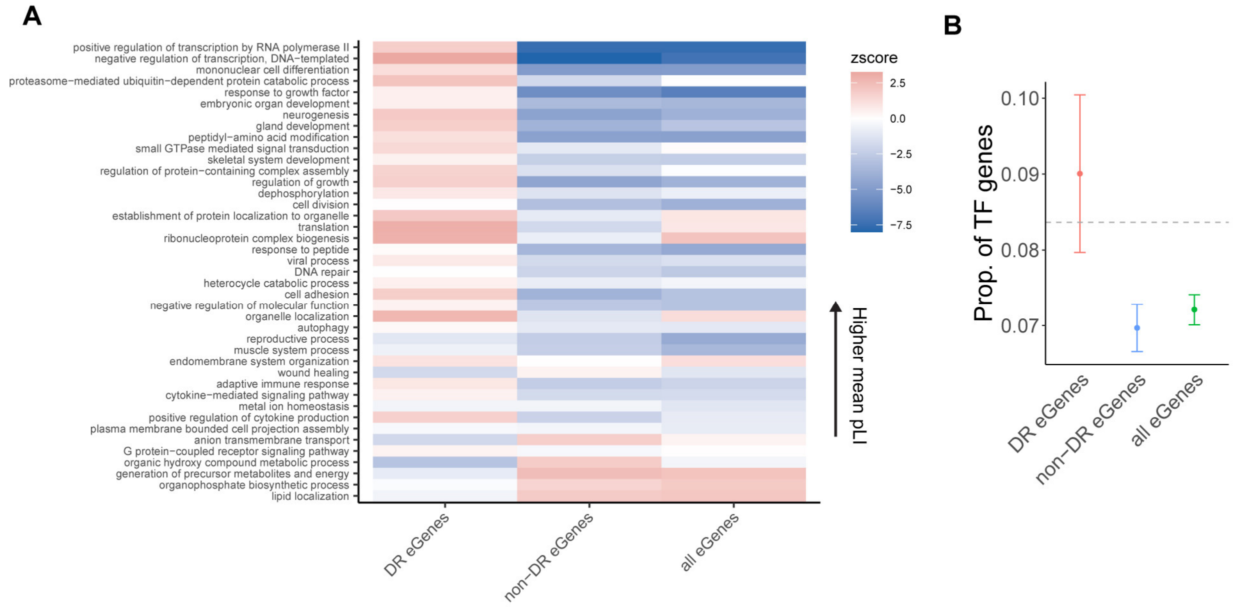


**Figure S16. Enrichment of eGenes in functional genes.**

(**A**) Enrichment of different group of eGenes (DR eGenes, non-DR eGenes, and all eGenes) among genes in 41 broadly defined Gene Ontology (GO) categories. The GO categories (y-axis) are sorted based on the average pLI value of the corresponding genes within each category. The color map represents enrichment (red) or depletion (blue) by enrichment Z scores.

(**B**) Proportion of transcription factor (TF) genes among DR eGenes (red), nonDR eGenes (blue), and all eGenes (green), respectively. The horizontal dashed line depicts the average fraction of TF genes in all genes that were tested in eQTL mapping. Error bars represent standard deviations.

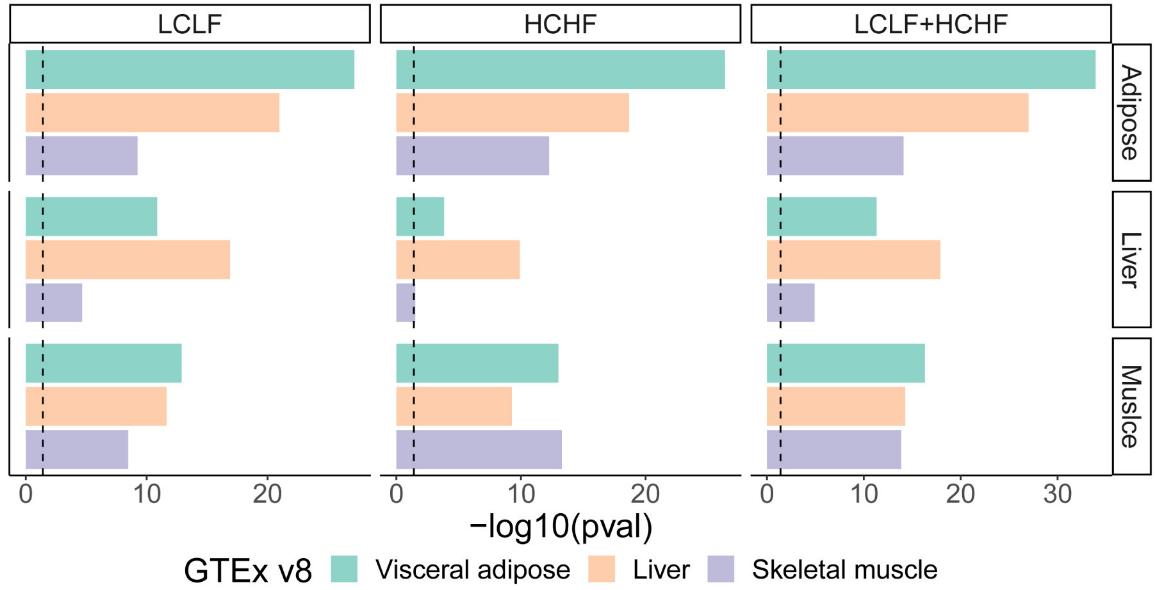


**Figure S17. Enrichment of baboon eGenes among human eGenes from GTEx.**

Enrichment of baboon eGenes (FDR<0.05) identified from each tissue (row) and diet (column) combination among human eGenes (q-value<0.05) in visceral adipose (green), liver (orange), and skeletal muscle (purple) from GTEx v8. Vertical dashed lines indicate a p-value threshold of 0.05.

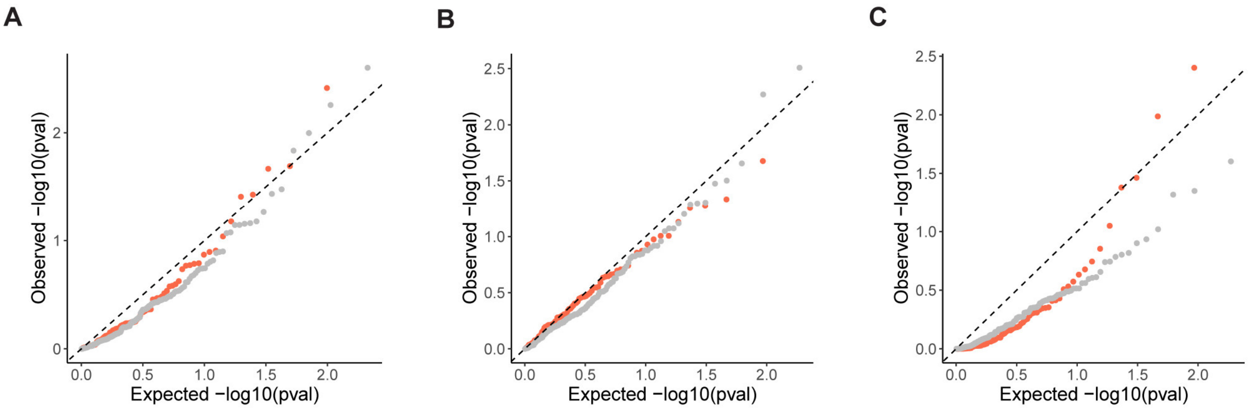


**Figure S18. Enrichment of baboon genes in human trait- or disease-associated genes from GWAS.**

Orange dots correspond to p-values of enrichment tests for metabolic-relevant traits relative to uniformly distributed p-values (dashed line). Gray dots correspond to p-values of enrichment tests for non-metabolic traits. Q-Q plot for enrichment of (**A**) baboon DR genes from DE analysis, (**B**) baboon genes that have similar expression levels with DR eGenes but do not harbor DR eQTLs, and (**C**) baboon non-DR eGenes in human trait- or disease-associated genes from GWAS.

| | LCLF Diet | HCHF Diet |
|---|---|---|
| **Energy, kcal/g provided by** | | |
| Carbohydrate, % kcal | 67.3 | 40.1 |
| Protein, % kcal | 19 | 20.4 |
| Fat, % kcal | 13.7 | 39.5 |
| Cholesterol mg/kcal | 0.02 | 1.75 |
| **Nutrients, % of ration**† | | |
| Protein, % | 15.5 | 21 |
| Arginine, % | 0.85 | 1.19 |
| Cystine, % | 0.23 | 0.37 |
| Glycine, % | 0.63 | 0.86 |
| Histidine, % | 0.41 | 0.52 |
| Isoleucine, % | 0.65 | 0.89 |
| Leucine, % | 1.33 | 2.02 |
| Lysine, % | 0.74 | 0.98 |
| Methionine, % | 0.32 | 0.39 |
| Phenylalanine, % | 0.77 | 1.06 |
| Tyrosine, % | 0.53 | 0.75 |
| Threonine, % | 0.56 | 0.77 |
| Tryptophan, % | 0.19 | 0.22 |
| Valine, % | 0.83 | 1 |
| Serine, % | 0.8 | 1.16 |
| Aspartic acid, % | 1.46 | 2.09 |
| Glutamic acid, % | 3.71 | 5.08 |
| Alanine, % | 0.89 | 1.21 |
| Proline, % | 1.37 | 1.78 |
| Taurine, % | 0 | 0.01 |
| **Primary fat source** | Vegetable | Lard |
| Fat (ether extract), % | 5 | 18 |
| Fat (acid hydrolysis), % | 6.2 | 19.4 |
| **Fatty acid composition, %** | | |
| Total saturated fatty acids, % | 1.17 | 7.33 |
| Total monounsaturated fatty acids | 1.33 | 7.4 |
| C18:2 linoleic | 2.1 | 2.39 |
| C18:3 linolenic | 1.9 | 0.14 |
| Other omega-3 polyunsaturated fatty acids | 0.23 | 0.14 |
| **Fiber (crude), %** | 8.3 | 4 |
| Neutral detergent fiber, % | 22.7 | 14 |
| Acid detergent fiber, % | 10.2 | 5.6 |
| **"Nitrogen-Free Extract" (by difference), %** | 54.7 | 41.2 |
| Starch, % | 28.5 | 22.21 |
| | | (Continues) |

(Continued)

| | | |
|---|---|---|
| Glucose, % | 0.25 | 0 |
| Fructose, % | 0.27 | 0 |
| Sucrose, % | 1.73 | 2.08 |
| Lactose, % | 0.15 | 0 |
| **Total digestible nutrients, %** | 76 | 88.5 |
| **Gross energy, kcal/g** | 3.84 | 4.11 |
| **Minerals** | | |
| Ash, % | 6.1 | 5.8 |
| Calcium, % | 1 | 0.84 |
| Phosphorus, % | 0.7 | 0.49 |
| Phosphorus (non-phytate), % | 0.4 | 0.19 |
| Potassium, % | 0.8 | 0.8 |
| Magnesium, % | 0.21 | 0.18 |
| Sulfur, % | 0.21 | 0.22 |
| Sodium, % | 0.35 | 0.52 |
| Choline, % | 0.55 | 0.1 |
| Fluorine, ppm | 24.3 | 4.1 |
| Iron, ppm | 378 | 315 |
| Zinc, ppm | 131 | 132 |
| Manganese, ppm | 118 | 118 |
| Copper, ppm | 37 | 18 |
| Cobalt, ppm | 0.76 | 0.59 |
| Iodine, ppm | 1.54 | 1.42 |
| Chromium (added), ppm | 1.79 | 0.35 |
| Selenium, ppm | 0.37 | 0.41 |
| **Vitamins** | | |
| Carotene, ppm | 1.1 | 1.1 |
| Vitamin K (as menadione), ppm | 3.2 | 2.4 |
| Thiamin hydrochloride, ppm | 11 | 14 |
| Riboflavin, ppm | 12 | 7.3 |
| Niacin, ppm | 123 | 92 |
| Pantothenic acid, ppm | 61 | 50 |
| Choline chloride, ppm | 1200 | 1468 |
| Folic acid, ppm | 2.2 | 8.8 |
| Pyridoxine, ppm | 15 | 12.02 |
| Biotin, ppm | 0.2 | 0.2 |
| Vitamin B-12, mcg/kg | 33 | 40 |
| Vitamin A, IU/g | 40 | 35 |
| Vitamin D-3 (added), IU/g | 7 | 5.4 |
| Vitamin E, IU/kg | 49 | 90 |
| Ascorbic acid, ppm | 541 | 407.7 |

**Table S1. Composition of the LCLF and HCHF diets***

*Based on specifications of the manufacturer, LabDiet, PMI Nutrition International, St. Louis, MO, USA. Because nutrient composition of natural ingredients varies, analysis can differ accordingly.
†Nutrients expressed as percent of ration on an as-fed basis except where otherwise indicated. Moisture content is assumed to be 10.0% for the purpose of calculations.

**Table S2. Metadata of sample collection.** Available as an excel file online.
**Table S3. Summary statistics of DR genes from DE analysis.** Available as a csv file online.
**Table S4. Summary statistics of sex-biased DR genes.** Available as a csv file online.
**Table S5. Summary statistics of all eGene-SNP pairs.** Available as a text file online.
**Table S6. Summary statistics of all DR eQTL-gene pairs.** Available as a text file online.
**Table S7. Lipid biomarkers.** Available as an excel file online.
**Table S8. Human GWAS metabolic relevant traits.** Available as a text file online.
**Table S9. Human GWAS non-metabolic relevant traits.** Available as a text file online.
